# Social reprogramming in ants induces longevity-associated glia remodeling

**DOI:** 10.1101/2019.12.19.883181

**Authors:** Lihong Sheng, Emily J. Shields, Janko Gospocic, Karl M. Glastad, Puttachai Ratchasanmuang, Arjun Raj, Shawn Little, Roberto Bonasio

## Abstract

Healthy brain aging is of crucial societal importance; however, the mechanisms that regulate it are largely unknown. Social insects are an outstanding model to study the impact of environment and epigenetics on brain function and aging because workers and queens arise from the same genome but display profound differences in behavior and longevity. In *Harpegnathos saltator* ants, adult workers can transition to a queen-like phenotypic state called gamergate. This caste transition results in reprogramming of social behavior and lifespan extension. Whether these changes in brain function and physiology cause brain remodeling at the cellular level is not known. Using single-cell RNA-sequencing of *Harpegnathos* brains undergoing caste transition, we uncovered shifts in neuronal and glial populations. In particular, the conversion of workers into long-lived gamergates caused the expansion of ensheathing glia, which maintain brain health by phagocytosing damaged neuronal structures. These glia cells were lost during aging in normal workers but not in longer-lived gamergates. We observed similar caste- and age-associated differences in ensheathing glia in other Hymenoptera as well as *Drosophila melanogaster*. We propose that enhanced glia-based neuroprotection promotes healthy brain aging and contributes to the extended lifespan of the reproductive caste in social insects.

## INTRODUCTION

Age-associated cognitive decline has an immense impact on human societies and is caused by genetic, epigenetic, and environmental factors, whose interplay is poorly understood^1–3^. Social insects, including ants, provide a fascinating model to study the epigenetic regulation of brain function and longevity^4–6^. The division of labor typical of social insect colonies is based on the distinct physical and behavioral phenotypes of highly related individuals organized in separate social castes. These individuals carry out distinct specialized tasks such as reproduction, foraging, defense, and nest maintenance^7, 8^. A notable difference between social castes is their life expectancy: in most species reproductive queens live much longer—often up to an order of magnitude longer—than sterile workers from the same colony^9^.

Brains of queens and various types of workers differ at a molecular level in the genes they express^4, 10, 11^ and often also at a structural level in the overall volume and relative size of anatomical sub-structures, such as the mushroom body^12–14^. However, whether brains from different social castes have substantially different cellular compositions that might contribute to caste phenotype has not been investigated, partly because we still lack a comprehensive molecular description of the variety of cell types that constitute a social insect brain.

The ant *Harpegnathos saltator* offers a unique opportunity to study the epigenetic regulation of phenotypic plasticity^6, 15, 16^. While in most ant species social castes are permanently established during the larval stage, adult *Harpegnathos* workers can acquire a queen-like phenotype and become reproductive individuals called “gamergates”^17, 18^. This adult caste transition results in a dramatic switch in social behavior^19–22^ and an extensive molecular reprogramming of the brain with hundreds of differentially expressed genes^19, 23^. Remarkably, workers that become gamergates also acquire queen-like longevity, with a 5-fold increase in average lifespan from 7 months to 3 years^24^.

Previous studies reported macroscopic changes in the brain of gamergates compared to workers, including a volumetric shrinkage of the optic lobe^25^, suggesting that extensive brain remodeling accompanies caste reprogramming. However, the questions of which cell types are most affected by the transition and most likely to contribute to the caste-specific regulation of behavior and longevity has not been explored.

To answer this question, we performed high-throughput single-cell RNA sequencing in brains from workers and gamergates and comparative transcriptomics analyses in other social insects as well as in young and old *Drosophila* brains. Our findings reveal deeply conserved processes of brain remodeling that may (1) contribute to complex social behaviors, (2) underlie the behavioral transition from worker to gamergate, and (3) promote healthy brain aging and increased lifespan via the expansion of neuroprotective glia.

## RESULTS

### Transcriptional types of neurons and glia in a social insect brain

We performed single-cell RNA-seq using 10x Genomics on brains harvested from workers (*n =* 6) and gamergates (*n =* 5) 30 days after initiating the caste transition (**Fig. 1a**). At this time, most of the dueling interactions within the transitioning colony have ceased^19^ and the newly converted gamergates have started to lay eggs. Importantly, caste-specific changes in gene expression were already established at day 30 and were captured by our single-cell RNA-seq, as demonstrated by a strong correlation (*r_S_ =* 0.42, *P =* 0.0001) with the differentially expressed genes that we previously identified by bulk RNA-seq at a later stage (day 120) of the transition^19^ (**Fig. S1a**).

**Figure 1.**
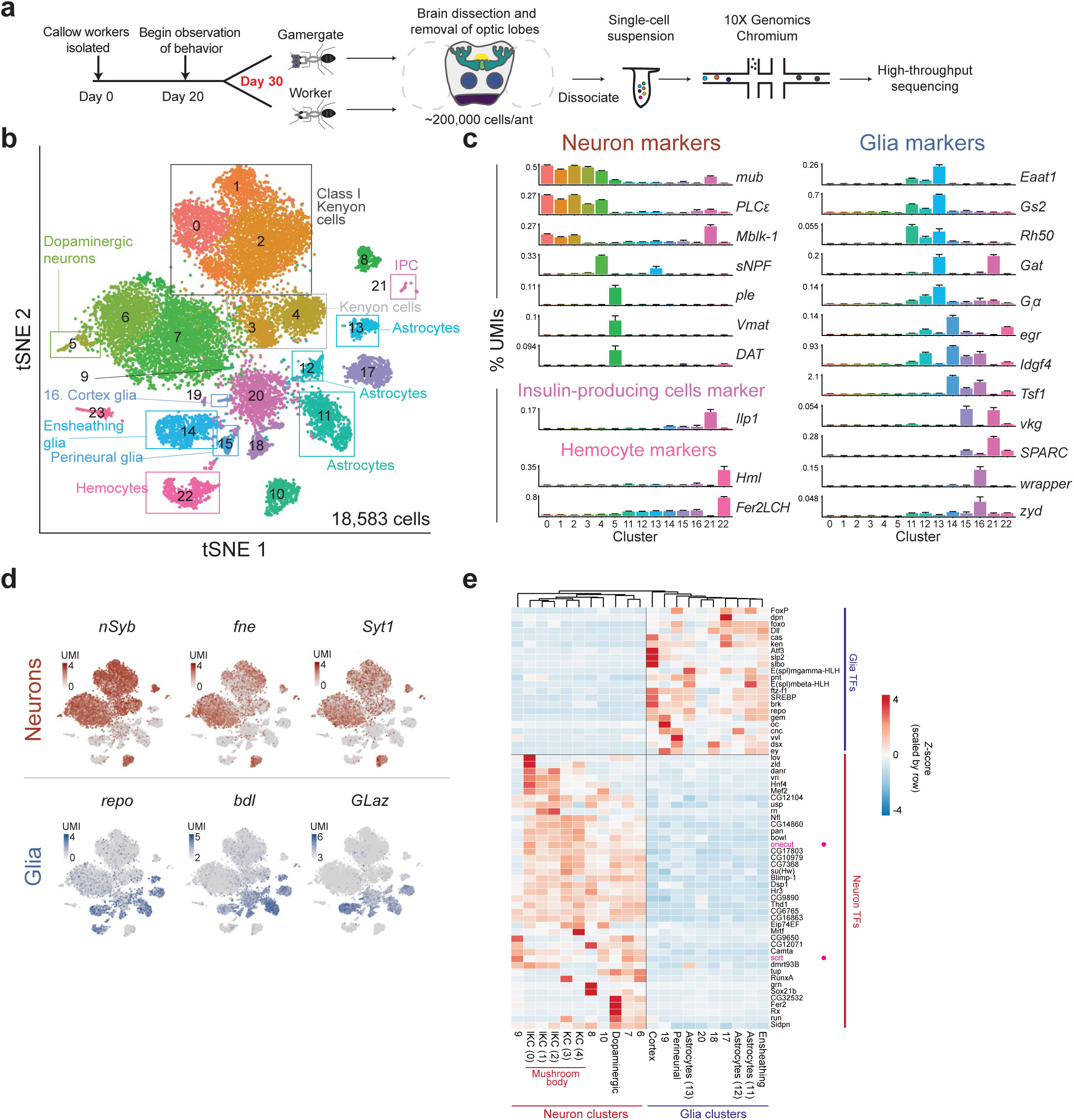
Single-cell transcriptomes from worker and gamergate brains. **a**, Scheme of the experiment. Workers and gamergates were separated on the basis of behavior and ovary status. Brains were dissected and optic lobes removed. The central brain, including mushroom bodies (dark green), ellipsoid bodies (green), fan-shaped bodies (yellow), and antennal lobes (blue), plus the gnathal ganglion (purple) were dissociated into a single-cell suspension and processed for single-cell RNA-seq. **b**, Annotated tSNE visualization of the clustering of 18,583 single-cell transcriptomes obtained by pooling all cells from 6 worker and 5 gamergate replicates. **c**, Selected marker genes for the clusters annotated in Fig. 1b. The *y*-axis shows the collapsed pseudo-bulk expression in each cluster (as % of total cluster UMIs) for the indicated gene. Bars represent the mean of 11 biological replicates + SEM. **d**, Heatmap plotted over global tSNE showing normalized UMIs per cell for known neuronal markers (red) and glia markers (blue). **e**, Heatmap for normalized expression levels (*z*-score) for the indicated genes in collapsed single-cell clusters. Only transcription factors with a |log_2_(neurons/glia)| > 1 are shown, but the columns were clustered on all transcription factors. Genes mentioned in the text as known neuronal transcription factors (pink) are indicated with circles.

To obtain a comprehensive description of cell types in the *Harpegnathos* brain, we first considered all samples, regardless of caste. We retained only cells with a minimum of 500 unique transcripts (as defined by unique molecular identifiers, UMIs) over at least 200 different genes and obtained 18,583 cells with a median of 987 UMIs per cell mapping to 11,926 genes and a UMI median of at least 815 for each biological replicate (*n =* 11, **Fig. S1b**). Using the Seurat pipeline^26^, we obtained 24 clusters (**Fig. 1b** and **Tables S1–3**), which we annotated using known markers from *Drosophila* (**Fig. 1c**).

Our clusters effectively separated neurons (0–10) and glia (11–20), confirming that we were able to capture characteristic transcriptomes of single cells (**Fig. 1d**). Neuron clusters were identified by the expression of previously defined markers *nSyb*, *fne*, and *Syt1*, whereas expression of known glia markers *repo*, *bdl*, and *GLaz* identified glia clusters^27–34^. Using additional established markers, we mapped cells from a majority of the clusters to corresponding types in the *Drosophila* brain (**Fig. 1c, Tables S1** and **S2**), including: Kenyon cells (KCs; *mub* and *PLCε*; clusters 0–4), dopaminergic neurons (*ple* and *Vmat*; cluster 5), three distinct clusters of astrocytes (*Eaat1, Gs2, Rh50*, *Gat*, and/or *G_i_α*; clusters 11–13), ensheathing glia (*egr, Tsf1*, and *Idgf4*; cluster 14), perineurial glia (*vkg* and *SPARC*; cluster 15), cortex glia (*wrapper* and *zyd*; cluster 16), insulin-producing cells (*Ilp1*; cluster 21), and hemocytes (*Hml* and *Fer2LCH*; cluster 22) .

Transcription factors are key specifiers of identity and function for all cells, including those in the brain^35–38^. Cell type-specific transcriptional networks have been observed in other single-cell experiments^28, 39^ and likely play a role in shaping the diversity and plasticity of cell types in the ant brain. We curated a panel of 423 transcription factors in the *Harpegnathos* genome based on their sequence conservation with *Drosophila* homologs (**Table S3**) and determined their expression pattern in single cells. Hierarchical clustering based on transcription factor expression alone separated neuron and glia clusters (**Fig. 1e**). This analysis confirmed the expression patterns of many known neuronal transcription factors including, for example, *onecut*^28^, which induces the transcription of neuron-specific genes^40, 41^, and *scrt*, which represses non-neuronal cell fates^42^.

Based on our clustering, 27% of the single cells recovered from *Harpegnathos* brains are glia (**Fig. S1c–d**). Although vast discrepancies have been reported in the relative abundance of glia when comparing histological and single-cell sequencing studies, all approaches point to a much larger glial compartment in mammals compared to *Drosophila* ^43–46^. Single-cell analyses that can be compared to ours recovered ∼25% of glia from human brains^47^ but only 5–10% from *Drosophila* brains^27, 28^ (**Fig. S1c–d**) or optic lobes^39^. Thus, glia in *Harpegnathos* are more abundant than in *Drosophila*, suggesting a prominent role for these crucially important cells^48^ in brain health and function in ants.

### Expanded mushroom bodies in the *Harpegnathos* brain

To better characterize neuronal populations in *Harpegnathos*, we re-clustered in isolation all the transcriptomes from neuronal clusters 0–10 (**Fig. 2a**, **Table S1** and **S4**), and found that the majority of neurons (54%, corresponding to 36% of all brain cells) expressed genes known to mark KCs in *Drosophila* (*mub*^49^ and *Pka-C1*^50^) and *Apis mellifera* (*PLCε* and *E75*^51–53^) (**Fig. 2b** and **S2a**). KCs are the principal component of mushroom bodies, the center of learning and memory in insects^54, 55^, which are known to be markedly large in social Hymenoptera^12, 56^, including ants^57^. Our single-cell RNA-seq confirms this conclusion, as KCs comprise a much larger fraction (54%) of neurons in *Harpegnathos* than in *Drosophila* (5–10%) (**Fig. 2c–d** and **S2b**), even after accounting for differences in the datasets by equalizing read numbers and UMI distributions (**Fig. S2c–f**). Immunofluorescence stainings for the KC marker Pka-C1 in *Harpegnathos* labeled structures with the anatomical features of mushroom bodies including a thick pedunculus (**Fig. 2e**, gray arrowhead) and prominent double cup-shaped calyces (**Fig. 2e**, white arrowhead), characteristic of Hymenoptera^12, 58^. Consistent with the increased frequency of KCs in our single-cell data, *Harpegnathos* mushroom bodies appeared to occupy a larger relative volume in the ant brain as compared to the corresponding structures in *Drosophila* (**Fig. 2e**). Along the same lines, western blots for Pka-C1 from total brain extracts revealed higher levels of this protein in ants as compared to flies (**Fig. 2f**).

**Figure 2.**
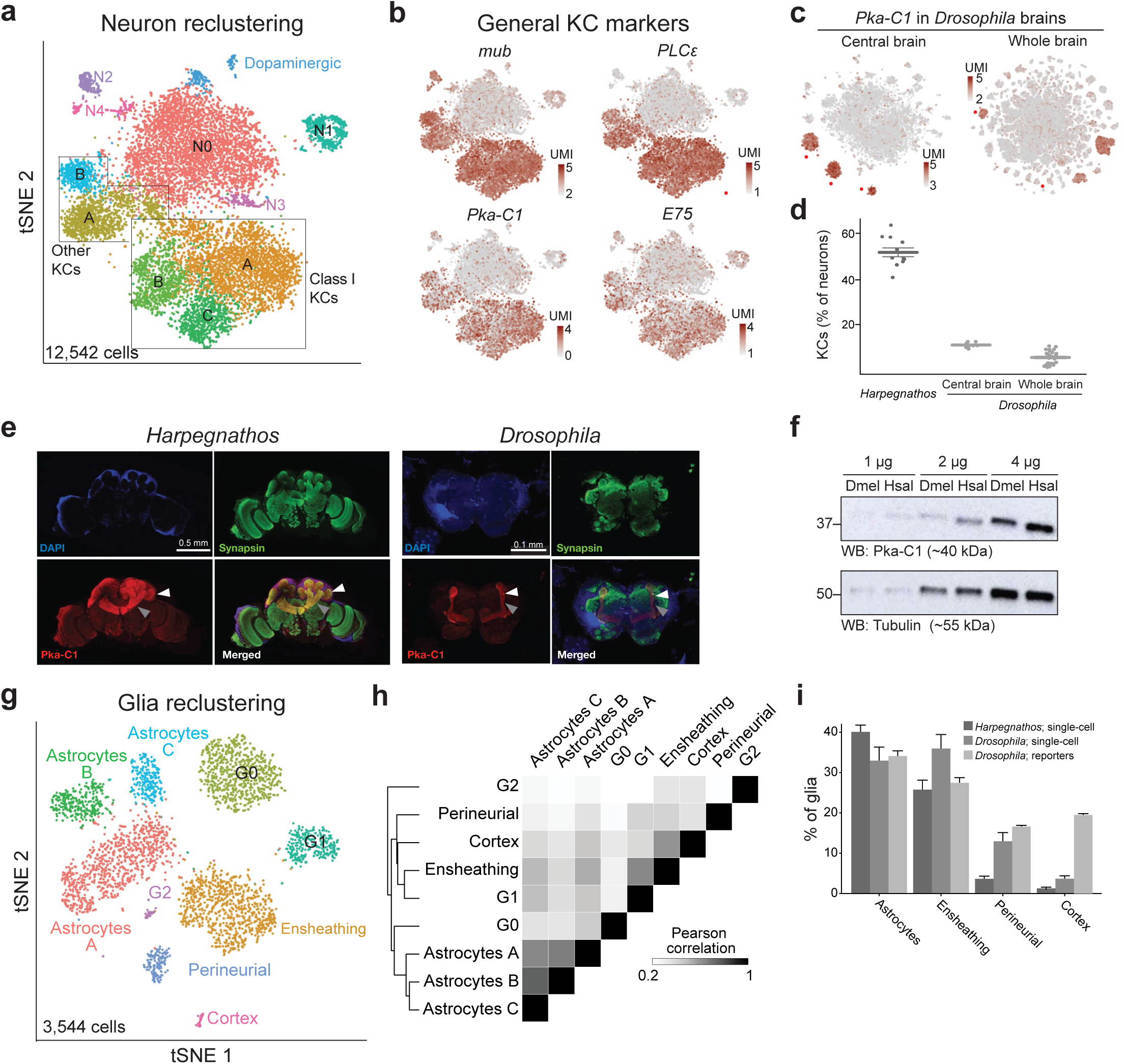
Distinctive features of *Harpegnathos* neurons and glia. **a**, Annotated tSNE visualization for the reclustering of neurons from 6 worker and 5 gamergate replicates at day 30. **b**, Heatmap plotted over neuronal tSNE showing normalized UMIs for known mushroom body markers from *Drosophila* (left) and *Apis mellifera* (right). **c**, Heatmap plotted over neuronal tSNE for the KC marker *Pka-C1* from two *Drosophila* single-cell RNA-seq datasets, one from the central brain after removing optic lobes (left; Croset et al. 2018) and one from the whole brain, inclusive of the optic lobes (right; Davie et al. 2018). **d**, Relative abundance of KCs as determined by percentage of neurons in clusters that express *Pka-C1* in *Harpegnathos* brains and in the two *Drosophila* single-cell RNA-seq datasets. Horizontal bars indicate mean ± SEM. **e**, Immunofluorescence for the neuronal marker synapsin and the KC marker Pka-C1 in Harpegnathos (left) and Drosophila (right) with DAPI as nuclear counterstain. Gray arrowhead, pedunculus; white arrowhead, calyx. **f**, Western blot for Pka-C1 in the indicated amount of total protein extract from *Drosophila* (Dmel) or *Harpegnathos* (Hsal) brains. Tubulin was used as loading control. **g**, Annotated tSNE visualization for the reclustering of glia from 6 worker and 5 gamergate replicates at day 30. **h**, Clustered heatmap showing the pairwise Pearson correlation score between collapsed transcriptomes (pseudo-bulk analysis) of glia clusters, considering only variable genes that were utilized to define the clusters by Seurat. **i**, Relative abundance of key glia subsets in *Harpegnathos* single-cell RNA-seq, *Drosophila* single-cell RNA-seq from the whole brain (Davie et al. 2018), and as defined by glial subset-specific expression of GAL4 (Kremer et al. 2017). Bars indicate means + SEM.

The larger mushroom body of *Harpegnathos* comprised a diverse repertoire of KC transcriptomes, as they separated into five clusters compared to the three clusters found in *Drosophila* (**Fig. 2a, c**). KCs were first described in the honey bee *Apis mellifera*^59^, where they are divided in three classes (I–III) based on their connectivity^12^. This classification appears to hold and translate to distinct transcriptional types in the *Harpegnathos* brain, as cells expressing known markers for class I KCs (IKC) *IP_3_R* and *Mblk-*1^60, 61^ clustered apart from other KCs (**Fig. S2g**). Gene ontology (GO) terms classically associated with mushroom body function were selectively enriched in transcriptionally distinct subtypes of IKCs, including “olfactory learning” (all IKCs), “learning or memory” (lKC A), and “associative learning” (lKC B) (**Fig. S2h**) suggesting functional sub-specialization of KCs in the *Harpegnathos* brain.

Thus, as typical of social insects^12, 56, 57^, the *Harpegnathos* brain contains an expanded mushroom body. In addition, our data revealed a transcriptionally diverse repertoire of KCs, perhaps conferring advanced learning and memory skills to navigate life in complex ant societies.

### *Harpegnathos* glia cells are diverse and include multiple subsets of astrocytes

Given the relative abundance of glia in *Harpegnathos* compared to *Drosophila* (**Fig. S1d**), we decided to analyze these cells further. We re-clustered and assigned identities to glia cells using genes homologous to known *Drosophila* markers (**Fig. 2g**, **S3a–c, Table S1** and **S5**). We identified cortex glia (*wrapper* and *zyd*)^28, 39, 62^, two clusters of astrocytes (astrocytes A and B in **Fig. 2g**; *Eaat1* and *Gat* or *Rh50*)^27, 28, 63–66^, ensheathing glia (*egr*, *Tsf1*, and *Idgf4*;^28^), and perineurial glia (*vkg* and *Tret*;^67–69^). An additional cluster (astrocytes C) displayed weak *Eaat1* expression and a transcriptome closely related to those of the two other types of astrocytes (**Fig. 2h**). Additionally, three clusters (G0, G1, and G2) expressed multiple glia markers but could not be unequivocally assigned to known transcriptional types, suggesting the existence of glia subtypes in *Harpegnathos* ants that are not present or have not yet been described in the central brain of *Drosophila* (**Fig. 2g**, **Table S5**).

The three distinct classes of *Harpegnathos* astrocytes comprised ∼40% of total glia cells, which was comparable to the frequency of astrocytes in *Drosophila* (**Fig. 2i**), as measured by fluorescence microscopy using genetic reporters^46^ and single-cell RNA-seq^28^. Similarly, the proportion of ensheathing glia in *Harpegnathos* (24% of all glia) was comparable to their frequency in genetically labeled *Drosophila* brains (**Fig. 2i**), suggesting that single-cell RNA-seq protocols capture efficiently the transcriptomes of these two cell types. On the other hand, perineurial and cortex glia were present at different frequencies across datasets. Perineurial glia, which envelope the outer surface of the brain and provide blood-brain barrier functions in insects were a three-fold larger fraction of *Drosophila* (15%) than *Harpegnathos* (4%) glia (**Fig. 2i**). This difference might in part be due to the larger ant brain having a lower surface-to-volume ratio, but we cannot exclude technical issues specific to *Harpegnathos* in recovering these cells. Indeed, technical difficulties seemingly affected the recovery of cortex glia in previous *Drosophila* single-cell RNA-seq studies, as evidenced by the much smaller fraction recovered with this technology (3.7%) compared to imaging-based estimates (19%)^46^. The former number is comparable to our quantifications by single-cell RNA-seq in *Harpegnathos* (**Fig. 2i**) and suggests that the honeycomb-like structure of this glia subtype^46^ might hamper their recovery in single-cell suspensions in both species.

The separation of *Harpegnathos* astrocytes into three clusters is a notable contrast with previous studies of the *Drosophila* brain^28^, in which astrocytes transcriptomes formed a single cluster (**Fig. S3d**). This distinction was independent of sequencing depth, as equalizing the number of glia cells and UMI distributions across datasets did not cause the three separate *Harpegnathos* clusters to merge (**Fig. S3d–e**). Astrocyte diversity has not been explored in insects, but there are reports of regional and functional heterogeneity among mammalian astrocytes^70–74^, which have been corroborated by transcriptomic analyses^75, 76^. Differences in marker genes (**Table S1** and **S5**) and associated enriched GO terms (**Fig. S3f**) for the three *Harpegnathos* clusters of astrocyte-like cells suggest that a similar functional heterogeneity of astrocytes might exist in ants.

Our results show that *Harpegnathos* brains contain a large fraction of glia cells, many of which belong to glia types described in *Drosophila*, while others display molecular fingerprints only found in ants, consistent with the possibility of functional specialization.

### Caste-specific brain remodeling includes the expansion of ensheathing glia in gamergates

In social insects, worker and queen brains are functionally and structurally distinguishable, with visible anatomical differences between the castes, especially in the mushroom body^12–14, 77^. In *Harpegnathos,* bulk transcriptional differences accompany behavioral and reproductive changes in workers that transition to gamergate status^19^. The occurrence of extensive brain remodeling during the caste transition is supported by morphometric analyses that show a loss in brain volume concentrated in the optic lobes^25^, but whether changes in cell composition correspond to this molecular and anatomical differences in the brain is completely unexplored.

We separated and compared single-cell transcriptomes from brains of workers (*n =* 6) and gamergates (*n =* 5) after 30 days of transition. The clustering profiles were broadly similar, demonstrating the absence of major batch effects (**Fig. S4a**); however, KCs type A, ensheathing glia, and perineurial glia displayed significant caste-specific changes in relative numbers (**Fig. 3a–b, S4b**). KC A neurons were relatively more abundant in gamergates than in workers (**Fig. 3a–b, S4c**) and were distinguishable from other KCs by their strong and specific expression of sNPF (**Fig. S4d**), which has been associated with the positive regulation of insulin signaling, lifespan, and other caste-specific traits^78–80^. KC A neurons also expressed *Sema2a* (**Fig. S4d**), a gene involved in axon guidance and remodeling^81^, suggesting gamergate-specific rewiring of neuronal connections involving sNPF-producing KCs in the mushroom body.

**Figure 3.**
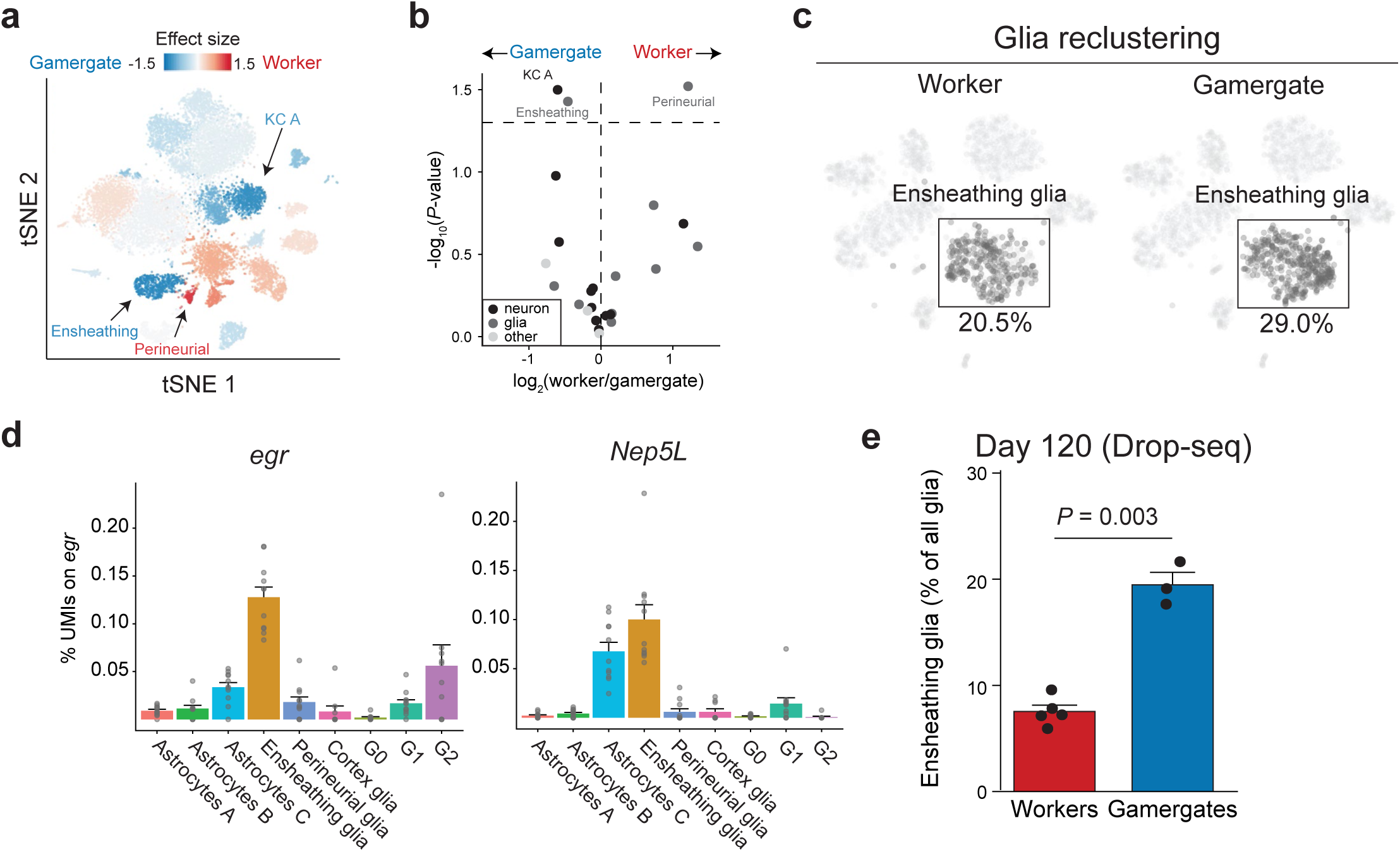
Cellular remodeling in the *Harpegnathos* brain after the caste transition. **a**, Heatmap plotted over global tSNE showing the changes in cell type abundance in workers vs. gamergate brains. The color scale indicates the effect size (mean/standard deviation). The three clusters that are significantly (*P*-value < 0.05, two-tailed Student’s *t*-test) affected are indicated. **b**, Volcano plot of log_2_(ratio) and -log_10_(*P*-value from a two-tailed Student’s *t-*test) for the relative frequency of each cluster in workers *vs.* gamergates. **c**, Visualization and quantification of worker (left) and gamergate (right) contributions to the ensheathing glia cluster in the reclustered tSNE for glia only (see Fig. 2g). Worker and gamergate datasets were downsampled to include the same number of total cells for comparison. **d**, Expression levels (% of cluster UMIs) across glia subsets for *egr* and *Nep5L*. Bars indicate means + SEM. **e**, Relative abundance of ensheathing glia as percentage of total glia in 120-day-old gamergates and workers as determined by Drop-seq. The *P*-value is from a two-tailed Student’s *t*-test.

Unexpectedly, caste-specific changes in glia composition were even more pronounced. The transition from worker to gamergate status resulted in a 64% reduction in perineurial (“surface”) glia (**Fig. S4e**) and a 41% expansion of ensheathing (“wrapping”) glia (**Fig. 3a–c**). The latter observation was of particular interest because, in *Drosophila*, ensheathing glia cells are essential to brain health and homeostasis as they remove damaged neuronal structures generated by stress, injury, or programmed cell death^48, 82^.

The expression of the phagocytic receptor Draper (*drpr*), its associated adaptor protein *dCed-6*, and the tumor necrosis factor (*Tnf*) homolog *eiger* (*egr*) are required for the phagocytic function of ensheathing glia in *Drosophila*^82–85^. *Drpr* knockout leads to neurodegeneration^86, 87^ as well as reduced lifespan in a *Drosophila* model of Alzheimer’s disease^88^, supporting the notion that ensheathing glia have a neuroprotective function. The *Harpegnathos* glia population that expanded during the worker–gamergate transition expressed *egr* and *Ced6*, confirming their phagocytic nature (**Fig. 3d, S4f**). Moreover, *Harpegnathos* ensheathing glia expressed the highest levels of a Hymenoptera-specific member of the neprilysin family, which we called *neprilysin-5-like* (*Nep5L*) (**Fig. 3d, S4g**). This family of metalloendopeptidases has been linked to the clearing of amyloid plaques in various experimental models^89, 90^.

As ensheathing glia are primarily responsible for the removal of dead or damaged neurons^82^; we wondered whether their expansion during the worker–gamergates transition was an acute and transient response to extensive brain remodeling. If that were the case, we reasoned that at a much later time point, after the gamergates have settled into their new social status, the numbers of ensheathing glia in their brain should return to control levels. Using Drop-seq^91^, we measured the relative abundance of ensheathing glia in older workers (*n =* 5) and gamergates (*n =* 3) at 120 days after the onset of caste transition. At this time point, the social hierarchy is fully established and the dueling interactions that lead to the social transition have long ceased^19^; nonetheless, the ensheathing glia component remained much larger (250%) in gamergates than in workers (**Fig. 3e**).

Thus, in just 30 days, the adult caste transition results in major plastic changes in cell composition of the *Harpegnathos* brain with an expansion of sNPF-producing KCs and neuroprotective ensheathing glia in individuals that acquire the queen-like gamergate phenotype. Furthermore, the changes in ensheathing glia persist well after the transition is completed, indicating a permanent, caste-specific expansion of this glia population.

### Upregulation of ensheathing glia markers in the reproductive caste across Hymenoptera

To determine whether the caste-specific expansion of ensheathing glia observed in *Harpegnathos* was conserved with other social insects we reanalyzed available RNA-seq data from an additional ant species with a worker–gamergate system, *Dinoponera quadriceps*, a carpenter ant with a more conventional social structure, *Camponotus planatus*, and the primitively eusocial wasp *Polistes canadensis*. For all three species, extensively replicated bulk RNA-seq data obtained from brains was available^92, 93^. We reasoned that changes in mRNA levels for genes expressed predominantly in ensheathing glia might be detectable even in bulk brain RNA.

Expression of three prominent and conserved ensheathing glia makers, *Tsf1*, *Idgf4*, and *Prx2540-1*, was consistently upregulated in the brains of reproductive individuals from all three species (**Fig. S5a**). Furthermore, an unbiased comparison of the top 25 genes most specifically expressed by *Harpegnathos* ensheathing glia (**Table S2**, cluster 14) revealed a clear trend toward higher expression in the reproductive castes in *D. quadriceps*, *C. planatus*, and *P. canadensis* (**Fig. S5b**). This trend was maintained when considering all 213 *Harpegnathos* ensheathing glia makers (**Table S2**): in all three species the proportion of these markers upregulated in reproductive individuals was significantly (P < 0.05 by Fisher’s exact tests) increased when compared to all genes (**Fig. S5c**).

Together, our comparative transcriptomic analyses indicate that an increased abundance of ensheathing glia is a conserved characteristic of the brains of the reproductive caste in social Hymenoptera.

### *Harpegnathos* ensheathing glia respond to brain injury

Glia cells provide essential support to brain function and are required for damage control in response to injury in both mammals and insects^48, 94^. In the mammalian brain, phagocytic microglia cells proliferate and migrate to sites of injury and are involved in clearing damaged tissue, modulating inflammation, and promoting remyelination in concert with oligodendrocyte precursors^95–98^. In *Drosophila* brains, these phagocytic and repair functions are carried out by ensheathing glia cells, which combine features of microglia and oligodendrocytes and activate conserved transcriptional networks and molecular pathways following injury^82, 99–101^.

*Harpegnathos* ensheathing glia expressed *egr* and *ced-6* (**Fig. 3d, S4f**), which are involved in the injury response in *Drosophila*; thus, we hypothesized that these cells would also respond to injury in *Harpegnathos*. We isolated young workers from stable, non-transitioning colonies and induced damage in their brains by puncturing them with a needle (**Fig. 4a, S6a**), which, in *Drosophila*, results in glia activation and proliferation at the injury site^85^. We performed Drop-seq^91^ on dissected brains one and three days after injury (median UMI = 448, **Fig. S6b**) and analyzed the glial compartment (**Fig. 4b**, **Table S1** and **S6**). Most clusters recovered in the Drop-seq experiment mapped to the same cell types identified in the deeper 10x Genomics dataset obtained for the day 30 worker–gamergate comparison (**Fig. S6c**).

**Figure 4.**
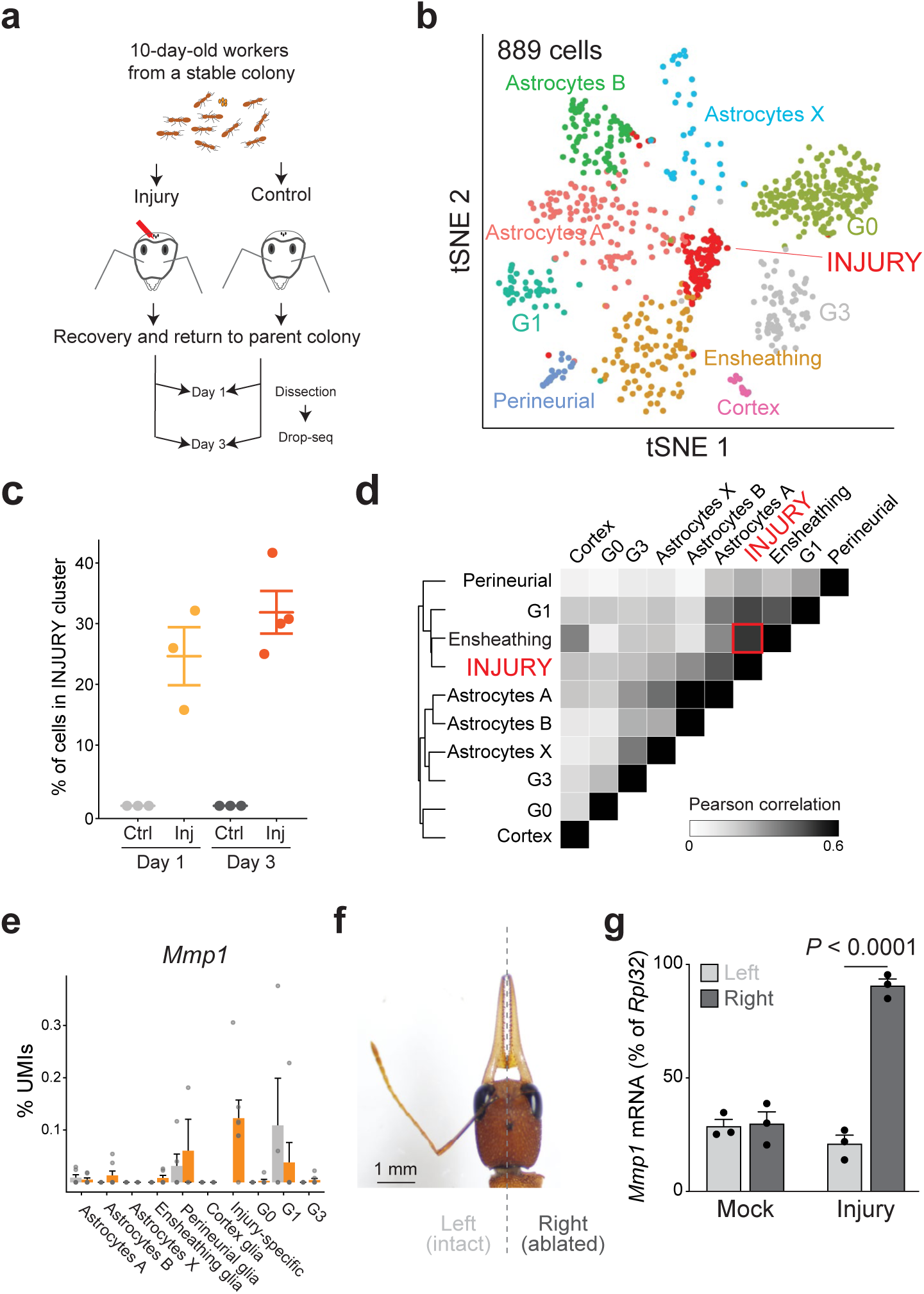
Brain injury causes activation of ensheathing glia. **a**, Scheme of the experiment. Needle-stabbed brains and age-matched control were analyzed by Drop-seq at day 1 and day 3 after injury. **b**, Annotated tSNE visualization of glia-only reclustering from pooled control and injury samples at day 1 and day 3 (*n =* 3 control and injury for day 1; *n =* 3 control and 4 injury for day 3). The injury-specific cluster is indicated. **c**, Relative frequency of cells in the injury-specific cluster as % of glia. Horizontal lines indicate means ± SEM. **d**, Clustered heatmap showing the pairwise Pearson correlation score between collapsed transcriptomes of glia clusters considering only variable genes that were utilized to define the clusters by Seurat. The high transcriptome-wide correlation between the injury-specific cluster and resting ensheathing glia is indicated by a red square. **e**, Expression levels (as % of cluster UMIs) for the activation marker *Mmp1*. Bars indicate means + SEM. **f**, Head of a 30-day-old *Harpegnathos* worker subjected to antenna amputation. **g**, RT-qPCR for *Mmp1* mRNA in the left (light gray) or right (dark gray) brain hemisphere after antennal ablation. Bars represent mean + SEM. *P*-value is from one-way ANOVA and Holm-Sidak test.

Interestingly, one subset of glia recovered in this experiment was unique to the injured brains and accounted for one third of all glia cells three days after injury (**Fig. 4b–c** and **S6d**). The transcriptomes from cells in this injury-specific cluster were closely related to the transcriptomes of resting ensheathing glia from uninjured brains (**Fig. 4d**). They expressed ensheathing glia markers *egr* and *Nep5L*, as well as the phagocytosis gene *ced-6* (**Fig. S6e**). These injury-specific ensheathing glia cells showed high expression of *Mmp1* (**Fig. 4e**), a known member of the response to injury in *Drosophila* that acts downstream of the phagocytic receptor *drpr*^102^ and is activated by amyloid plaques^88^.

To confirm that the observed transcriptional response was occurring as a local response to brain injury, we utilized a different experimental paradigm, whereby surgical amputation of an antenna (**Fig. 4f**) results in axonal degeneration and localized damage in the antennal lobe, which is known to induce a local glial response in *Drosophila*^85, 103^. Antennal ablation caused the upregulation of the activated glia activation marker *Mmp1* in the damaged but not in the contralateral hemisphere (**Fig. 4g**). No difference between right and left antennal lobes was observed in mock-treated animals.

Together, these observations suggest that, similar to their function in *Drosophila*, ensheathing glia respond to brain damage in *Harpegnathos* by activating the injury response, which results in a distinct transcriptional profile, including the upregulation of *Mmp1*.

### Gamergates are resistant to aging-associated decline in ensheathing glia

A key difference between the castes of social insects is that queens live much longer (often 10-fold or more) than workers from the same species^9^. Even more remarkably, the lifespan of a *Harpegnathos* individual is dramatically affected by changes in its social status, as the worker– gamergate transition results in a 5-fold increase in lifespan^24^. Given the importance of glia in protecting brains from the insults of time and disease^104–107^, we hypothesized that an expanded population of ensheathing glia in gamergates might contribute to their longer life expectancy and that the accelerated aging trajectory of workers might be accompanied by a change in the opposite direction.

Using Drop-seq, we profiled brains from 5-day-old pre-transition individuals, as well as 30-, 90-, and 120-day-old workers (*N* ≥ 3 per time point), corresponding to 2.4%, 14%, 43%, and 57% of their average 210 days lifespan^24^. The number and transcriptional identity of the glia clusters in these experiments were similar to those observed in the 10x Genomics profiling and injury experiments above (**Fig. S7a–d**). Ensheathing glia experienced a drastic decline during aging in *Harpegnathos* workers, especially between day 30 and day 90 and they were at their lowest numbers in 120-day-old workers (**Fig. 5a–b** and **S7e**). Notably, brains of 120-day-old gamergates contained significantly more (2.5-fold increase, *P =* 0.003) ensheathing glia cells (**Fig. 5b**, blue squares).

**Figure 5.**
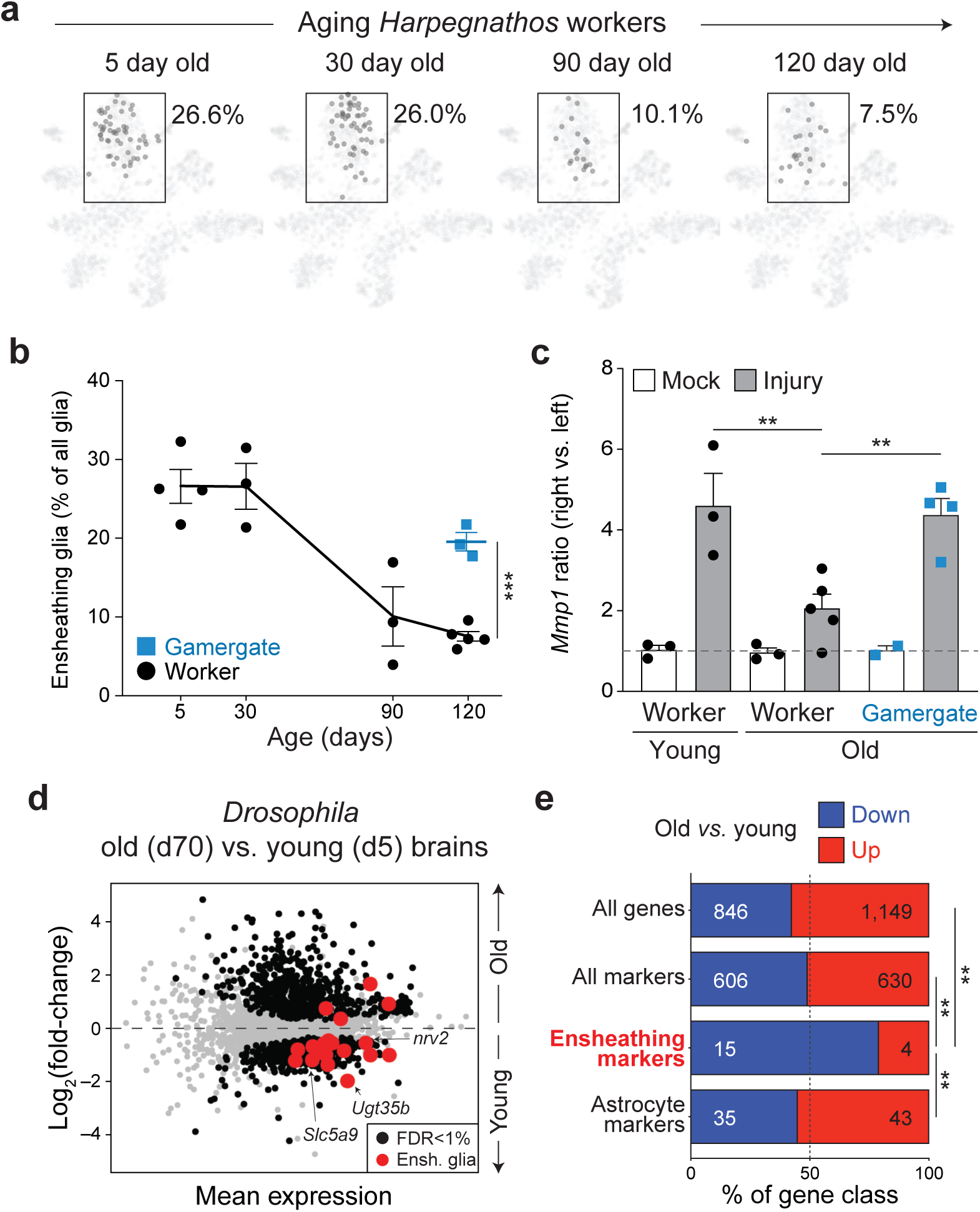
Ensheathing glia decline during aging in *Harpegnathos* and *Drosophila*. **a**, Visualization and quantification of the contributions from workers of different ages to the ensheathing glia cluster in the Drop-seq t-SNE. Cells from each time point were downsampled to the same total cell number for comparison. **b**, Dynamics of the ensheathing glia cluster during aging. Each point represents a biological replicate. The line connects the means and error bars indicate ± SEM. The mean for the gamergate sample is shown as a thicker bar. ***, *P* < 0.001 from a two-tailed Student’s *t*-test. **c**, Ratio of *Mmp1* mRNA (RT-qPCR) in the right vs. left brain hemisphere in ants subjected to ablation of the right antenna (gray) or a mock treatment as control (white). Activation of ensheathing glia as measured by upregulation of *Mmp1* was compared in young (30 day-old, left) and old (120–150 day-old, right) individuals. Bars represent mean + SEM. The dashed line indicates a ratio of 1, i.e. no difference between right and left hemisphere. **, *P*-value < 0.01 from one-way ANOVA and Holm-Sidak test. **d**, MA plot of RNA-seq data from brains of young (day 5, *n =* 3) and old (day 70, *n =* 4) *Drosophila* females. All 4,436 genes identified as cell type-specific markers by Davie et al.^28^ are shown. Differentially expressed (FDR < 1%) genes are shown in black. Differentially expressed (FDR < 1%) ensheathing glia marker genes are in red. **e**, Proportion of indicated gene classes that were significantly upregulated (red) or downregulated (blue) in old vs. young brains according to RNA-seq. **, *P*-value < 0.01 from one-sided Fisher’s test.

Consistent with the decline in ensheathing glia numbers in their brains, old workers exhibited a weaker transcriptional response to neuronal damage compared to young workers, as shown by a decreased upregulation of the glia activation marker *Mmp1* (**Fig. 5c**). Importantly, upon antenna ablation, old gamergates showed a much stronger *Mmp1* upregulation than age-matched old workers (**Fig. 5c**), indicating that the increased numbers of ensheathing glia in gamergates retain the ability to respond to brain damage into old age.

To determine whether the decline in ensheathing glia is a conserved feature of aging brains, we analyzed the expression of ensheathing glia markers in brains of young (day 5) and old (day 70) *Drosophila* females. RT-qPCR for *Slc5a9* and *Slc5a8*, two genes highly specific for ensheathing glia in *Drosophila* (**Fig. S8a, c**) and homologous to the *CG6723* marker gene in *Harpegnathos*, showed a 3-fold downregulation in the whole brain (**Fig. S8b, d**). Genome-wide transcriptome analyses of brains from young and old flies confirmed and extended these observations as 15 ensheathing glia marker genes were significantly (FDR < 1%) downregulated in aged individuals compared to only 4 that were upregulated (**Fig. 5d**). This bias toward downregulation in aged brains was evident also when considering all ensheathing glia marker genes, regardless of statistical cutoffs (**Fig. S8e**). Furthermore, the decline in expression was specific to ensheathing glia genes, as no similar trends were observed when considering all genes, all genes identified as cell-type specific markers by single-cell RNA-seq^28^, or astrocyte marker genes (**Fig. 5e**).

Thus, the expansion of ensheathing glia in *Harpegnathos* gamergates counters a deeply conserved natural decline in the abundance of this cell type in aging brains and endows them with an enhanced ability to respond to brain damage at an older age.

## DISCUSSION

Social insects provide an exceptional opportunity to study how epigenetic pathways regulate brain function and longevity. While many studies, including ours, have reported differences in gene expression between brains from different castes in bulk^11, 19, 92, 108, 109^, a single-cell RNA-seq approach was needed to analyze cellular plasticity and identify cell types most likely to contribute to caste-specific traits. We leveraged the unique phenotypic plasticity of *Harpegnathos* combined with the high-quality genomic resources^23^ in this ant species to obtain a comprehensive single-cell representation of the brain of a social insect. We found larger populations of mushroom body neurons and glia cells than in *Drosophila* and, within them, two specific subpopulations of cells that expand significantly as workers switch caste to become reproductive gamergates, sNPF-expressing KCs, and ensheathing glia. The latter showed an age-dependent decline in workers that was greatly attenuated in gamergates, suggesting that an enlarged ensheathing glia compartment provides heightened neuroprotection in the aging brain of gamergates, contributing to their 5-fold longer lifespan.

### Neuronal and glia cell types in the ant brain

Despite the evolutionary distance between ants and flies (> 350 million years), we were able to assign identities to a majority of single-cell clusters from *Harpegnathos* brains using known markers and transcription factors expressed in corresponding cell types in *Drosophila* (**Fig. 1**). The major distinction among neurons was between two types of KCs, IKCs (three clusters), as previously defined in honey bees, and other KCs (two clusters), which together account for a remarkable 54% of all *Harpegnathos* neurons outside of the optic lobe (**Fig. 2**). The observation that social Hymenoptera have larger mushroom body is far from novel^12, 56^, but our single-cell profiling also revealed unexpected complexities in this large population of KCs, which mapped to five different clusters. The single-cell transcriptomes for these different subsets of KCs, including the one uniquely affected during the caste transition (see below) provide molecular entry points to investigate the behavioral functions of different KC subtypes in social insects.

In addition to an enlarged mushroom body, our single-cell RNA-seq analyses revealed a prominent glia compartment in *Harpegnathos* (27%) compared to 5–10% as reported by multiple single-cell RNA-seq studies in *Drosophila*^27, 28, 39^. We detected glia of known insect subtypes, including surface, cortex, and neuropile glia, the latter further divided into ensheathing glia and astrocyte-like cells. Three relatively small clusters (named G0, G1, and G2) did not express clearly identifiable markers and therefore constitute either ant-specific glial cells that do not exist in *Drosophila*, or cells that exist in *Drosophila* but that have not been captured by existing studies. Additional evidence for the complexity of ant glia was found in the presence of three astrocyte subsets (**Fig. 2**), suggesting a sub-specialization of this cell type in ants.

### Caste-specific differences

Among the most remarkable properties of social insects is the dramatic difference in the behavioral and physiological phenotypes that originate from the same genome and give rise to the different social castes. Differences in the relative size of the brain and its anatomical compartments between different castes and even between individuals from the same caste allocated to different tasks (e.g. nursing vs. foraging) have been reported^12–14, 110–113^, but a comprehensive molecular comparison of cell types across castes was lacking. Here, we performed this comparison by taking advantage of the unique ability of *Harpegnathos* workers to convert to queen-like gamergates. We found evidence of remarkable caste-specific cellular plasticity in cell type composition of both neurons and glia, including an increase in the relative abundance of sNPF-expressing mushroom body KCs and ensheathing glia in gamergates accompanied by a relative reduction in perineurial glia (**Fig. 3**).

The changes in ensheathing glia caught our attention because they were substantial (41% increase at day 30, 250% at day 120) and consistent across biological replicates, time points, and sequencing technology (10x Genomics and Drop-seq). Single-cell RNA-seq can only provide relative numbers, and therefore we cannot formally exclude that the apparent specific expansion of ensheathing glia in gamergates is the result of a simultaneous loss in absolute numbers of most of the other brain cell types, however the most parsimonious explanation is that ensheathing glia respond to caste reprogramming by increasing in numbers, most likely via proliferation. It is known that insect glia proliferates in response to injury^85, 100, 101, 114^, but to our knowledge this is the first observation of a programmed, regulated expansion of a glia subset in otherwise healthy brains. We detected caste-specific differences in the expression of a majority of ensheathing markers in three other social insects (**Fig. S5**), *D. quadriceps* and *P. canadensis*, both with a flexible worker caste, similar to *Harpegnathos*, and also in the ant *C. planatus*, which has a fixed social structure with developmentally defined workers and queens.

Together, these observations suggest that glia plasticity, and more specifically an increased frequency of ensheathing glia in reproductive individuals are a conserved caste-specific feature of social insect brains.

### Ensheathing glia in injury and aging

In *Drosophila*, ensheathing glia play a critical role in the response to brain injury. They phagocytose degenerated axons after nerve damage via a well-studied molecular pathway that involve the phagocytic receptor Drpr, its adaptor protein dCed-6, and a signaling pathway that include TRAF4, JNK, and STAT92E^115^. Two lines of evidence suggest that the analogous cell type in *Harpegnathos* retains this function. First, the cells that we identified as ensheathing glia expressed *ced-6* as well as *egr*, which are required for the response to brain injury in *Drosophila*^85^. Second, after acute brain damage in *Harpegnathos* workers, a new glial cluster appeared, composed of cells with a transcriptional profile closely similar to that of ensheathing glia in uninjured brains, but expressing the activation marker *Mmp1* (**Fig. 4**), which is downstream of *Drpr*^88^. It is noteworthy that the functional homology of *Harpegnathos* ensheathing glia with neuroprotective glia in other animals might extend beyond insects, as microglia, the major glia cell type that responds to injury in mammalian brains^98, 116, 117^, express *Tnf*, the mammalian homolog of the ensheathing glia marker *egr*.

In mammals, glia composition and properties display age-related dynamics^107^, including a shift to a more neuroprotective function as the brain ages^118^, and restoration of the phagocytic function of microglia in aged mice improves their cognitive performance^119^. In *Drosophila*, glia are thought to play a role in regulating health and lifespan^105^ and ensheathing glia from aged individuals are less effective in clearing accumulated neuronal debris^102^, suggesting that declining ensheathing glia might be a cause rather than a consequence of aging. Given the extreme difference in lifespan between workers and gamergates in *Harpegnathos,* we hypothesized that the glia composition from the different castes would display divergent kinetics during aging. In fact, aging workers exhibited a progressive and dramatic loss of ensheathing glia cells (**Fig. 5a–b**), whereas gamergates were able to maintain them at higher numbers over the same time span (**Fig. 3e, 5b**). Consistent with this, old gamergates retained the ability to mount a response to neuronal damage, which was lost in old workers (**Fig. 5c**). Finally, transcriptomic analyses in *Drosophila* revealed that ensheathing glia marker genes are downregulated in older brains (**Fig. 5d–e**), indicating that aging-associated decline of ensheathing glia-based neuroprotection is a deeply conserved phenomenon.

### Conclusions and outlook

Single-cell RNA-seq profiling of *Harpegnathos* brains as they undergo the adult caste transition from worker to gamergate revealed plastic events in neurons and glia. Neuroprotective ensheathing glia cells were substantially expanded in gamergates as early as day 30 of the transition and they were severely reduced in numbers in old workers but not age-matched gamergates. Based on these observations, we propose that a homeostatic expansion of ensheathing glia in *Harpegnathos* gamergates promotes healthy brain aging by increasing its repair capacity, thus contributing to a prolonged lifespan. Along similar lines, we speculate that ensheathing glia play a broader role in regulating lifespan differences between castes in other social insects and that elevated numbers and proper function of phagocytic glia are required for healthy brain aging across the animal kingdom.

## METHODS

### Worker–gamergate transitions

*Harpegnathos* colonies were housed in plastic boxes with a plaster nest chamber in a temperature- and humidity-controlled ant facility on a 12-hour light cycle and maintained on a live cricket diet as previously described^19^. To induce worker-gamergate transitions we transferred 20 callow females (3–4 day old; backgrounds DR-91, TL, DR-105, and DR-101) from mature colonies to a new nest with four males. Each ant was individually painted with a unique two-color combination. After 20 days, ants were monitored for signs of caste-specific behavior, including foraging activity to identify workers and egg-laying to identify for gamergates. Ten days later (for a total of 30 days of transition), individuals were sacrificed and brains were collected by dissection. The caste of each individual was confirmed by inspecting ovaries for the presence of mature oocytes. Only individuals who laid eggs and had mature oocytes in their ovaries by day 30 were classified as gamergates, whereas only individuals who foraged and had inactivated ovaries were considered workers.

### Aging workers and gamergates

Newly eclosed ants from four stable colonies (backgrounds DR-91, TL, DR-105, DR-101) were painted individually with a unique two-color combination and returned to the colony of origin. Brains were collected after 5, 30, 90, and 120 days. Although spontaneous conversion of workers to gamergates in stable colonies is rare, the ovaries of the collected individuals were inspected for the absence of mature oocytes to confirm worker status. 120-day-old gamergates were collected from stable colonies and their ovaries were inspected to confirm their gamergate status.

### Stabbing brain injury

Ten-day-old callow workers were separated from stable colonies and briefly anesthetized on dry ice. Each ant was punctured with a tin sterile insect pin (FST #26002-15) in the proximity of the ocelli. Controls were age-matched callow workers that were subjected to the same experimental procedure minus the injury. After thirty minutes of recovery, ants were returned to their colony. Brains were collected 1 or 3 days after injury.

### Antenna ablation and RT-qPCR

Thirty- and 120-day-old workers were collected from stable colonies and briefly anesthetized on dry ice. The right antenna was surgically removed at the base of the head using scissors heat-sterilized with a Germinator 500 (Braintree scientific). As a control, age-matched workers were subjected to the same experimental procedure, including anesthesia, but the antenna was not removed (mock treatment). Thirty minutes after amputation, the ants were returned to their colony. Brains were harvested three days after injury and the two hemispheres were separated. RNA was purified with TRIzol from each hemisphere and expression of *Mmp1* was quantified by RT-qPCR using Power SYBR Green RNA-to-CT (Thermo Fisher Scientific). *Rpl32* was used as a reference gene^19^. Primer sequences are listed in **Table S8**.

### Single-cell dissociation

For consistency with our previous study^19^, we removed the optic lobes during all brain dissections. To dissociate cells, 1 mL of papain solution (5 mL DMEM medium/vial with 45 µM actinomycin D to block new transcription during processing^120^) was added to each brain, which were rotated at room temperature for 12 minutes. Brains were washed twice with DMEM, once with PBS-B (PBS + 0.01% BSA), and resuspended in 200 µL PBS-B, pipetting up and down with 200 µL tips about 20 times and gel loading tips about 30 times on ice. Samples were incubated for 5 minutes at room temperature and pipetting was repeated. Samples were filtered on a 40 µm Flowmi strainer and live cells were counted by Trypan Blue exclusion using both a Countess II and a hemocytometer.

### 10x Genomics

For each biological replicate, 3–4 brains (cell viability > 80%) were pooled. Samples were mixed with *Drosophila* S2 cells at a 10:1 cell ratio as a quality control. Cells were added to the RT mix with the aim of capturing the transcriptomes of ∼5,000–10,000 cells. All downstream cDNA synthesis (12 PCR cycles), library preparation, and sequencing were carried out as instructed by the manufacturer (10x Genomics Chromium Single Cell 3’ Reagent Kit v2 User Guide RevD), with minor modifications. Custom barcodes were used as indexes. After PCR, libraries were sequenced on an Illumina NextSeq 500. The read configuration was 26 bp (read 1), 8 bp (index 1), and 58 bp (read 2). Libraries were loaded at 2.0 pM.

### Drop-seq (injury and aging)

For each sample two brains were pooled. Ant cells were mixed with *Drosophila* S2 cells at a 10:1 ratio as a quality control. The final cell suspension was diluted to 100 cells/µL in PBS and 1.5 mL of the cell suspension was loaded for each Drop-seq run. Barcoded beads were resuspended in freshly made lysis buffer, composed of 200 mM Tris-HCl (pH 8.0_RT_), 20 mM EDTA, 6% Ficoll PM-400 (GE Healthcare/Fisher Scientific), 0.2% sarkosyl (Sigma Aldrich), and 50 mM DTT, at a concentration of 120 beads/mL. The flow rate for cells and beads were set to 4,000 mL/h, while the droplet generation oil (Bio-rad) was flown at 15,000 mL/h. Droplets were generated and collected in a 50 mL Falcon tube for a run time of 15 minutes. Droplet breakage was performed by with Perfluoro-1-octanol (Sigma-Aldrich), followed by reverse transcription, exonuclease I treatment, and amplification of cDNA according to the Drop-seq protocol from the McCarroll lab (http://mccarrolllab.org/dropseq/), with minor modifications. cDNA was amplified using Terra PCR Direct Polymerase Mix Kit. After two rounds of purification with 0.6x SPRISelect beads, we tagmented 600 pg of DNA using the Nextera XT DNA Sample Preparation Kit (Illumina, cat # FC-131-1096). Libraries were further amplified with 12 PCR cycles using custom P5-TSO hybrid and custom Nextera-compatible primers with different indexes. Libraries were loaded at 2.0 pM. Custom read 1 and index 2 primers were used, and libraries were sequenced on an Illumina NextSeq500. The read configuration was 20 bp (read 1), 8 bp (index 1), 8bp (index 2), and 56 bp (read 2).

### Western blots

*Harpegnathos* and *Drosophila* brains were dissected, separated from the optic lobes, homogenized in protein extraction reagent (Thermo Fisher Scientific, #89822) with protease inhibitors, frozen in -80 °C for at least 30 minutes, and then thawed for 15 minutes at room temperature with gentle rocking. Lysates were centrifuged at 16,000g for 20 minutes to remove insoluble material. Antibodies used were: anti-tubulin: DSHB E7 (1: 500), anti Pka-C1: Cusabio (1:2000). Anti-mouse IgG (H+L): Jackson immune Research (1:5000), Anti-rabbit IgG (H+L): Jackson immune Research (1:5000).

### Immunofluorescence

*Harpegnathos* brains were dissected in DMEM medium and fixed in 4% paraformaldehyde overnight at 4 °C. Fly heads were dissected in 1x PBS and fixed in 4% paraformaldehyde for 2 hours at room temperature. Washed in PBS twice and embedded with 4% agarose gel in PBS. Thick sections (100 µm) were obtained with a Vibratome (Leica VT 1000S), permeabilized in PBST (1x PBS + 0.5% Triton X-100) two times for 10 min, and then blocked with 5% goat serum in PBST for at least 1 hour at room temperature. Sections were incubated with primary antibodies overnight at 4°C in 5% goat serum in PBST, washed in PBST three times for 20 min, incubated with appropriate fluorescently labeled secondary antibodies for 2 hours in PBST, washed 6 times with PBST and stained with DAPI for 10 min. Finally, samples were washed three times for 5 min then mounted in anti-fade mounting medium (Vector Laboratories, H-1000). Sections were imaged with a Leica SPE laser scanning confocal microscope. Antibodies used were: anti-Synapsin: DSHB 3C11 (1: 20), anti Pka-C1: Cusabio (1:1000). Anti-mouse IgG (H+L), Alexa Flour 488: Thermo Fisher Scientific (1:500), Anti-rabbit IgG (H+L), Alex Flour 568: Thermo Fisher Scientific (1:500).

### Gene expression in young and old *Drosophila* brains

For the aging experiment, *y1w\** flies were raised at 25°C and 50% humidity on a 12-hour light– dark cycle using standard Bloomington Drosophila Medium (Nutri-Fly). Newly eclosed flies, over a 24-hour period, were transferred into fresh food vials. After 48 hours, ten mated females were collected under mild CO_2_ anesthesia and transferred into fresh food vials. Flies were maintained in the same conditions as described above and transferred to new food vials twice a week. After 5 days (young) and 70 days (old), brains were dissected and RNA was purified with TRIzol. RNAs were quantified using the Power SYBR Green RNA-to-CT 1-step kit.

For library preparation, polyA+ RNA was purified using Dynabeads Oligo(dT)_25_ (Thermo Fisher) beads and constructed into strand-specific libraries using the dUTP method^121^. UTP-marked cDNA was end-repaired using end-repair mix (Enzymatics, MA), tailed with deoxyadenine using Klenow exo-(Enzymatics), and ligated to custom dual-indexed adapters with T4 DNA ligase (Enzymatics). Libraries were size-selected with SPRIselect beads (Beckman Coulter, CA) and quantified by qPCR before and after amplification. Sequencing was performed in paired-end mode on a NextSeq 500 (Illumina, CA).

### Single-cell data processing

Sequencing files from both 10x Genomics and Drop-seq data were processed using Drop-seq tools v2.0.0, developed by the McCarroll Lab (https://github.com/broadinstitute/Drop-seq). Sequencing data was aligned to a combined reference of the current *Harpegnathos saltator* assembly^23^ (Hsal_v8.5) and the *Drosophila melanogaster* genome assembly (assembly Release 6 plus ISO1 MIT). Reads mapping to features from the *Harpegnathos* annotation (Hsal_v8.5) were counted and a UMI matrix was produced for all cells with a minimum number of 150 genes.

### Single-cell data clustering and visualization

#### 10x Genomics data analysis

Analysis of the UMI matrix produced by Drop-seq tools was performed with Seurat v2.3.4^26^. Cells with more than 200 genes and 500 UMIs were retained, with each gene being detected in at least 3 cells. Cells from each sample were log-normalized with a scale factor of 10,000. Data were then scaled so the mean of each gene across cells was 0 and the variance across cells was 1. As the data was generated in three separate experiments (experiment 1: 2 workers and 1 gamergate, experiment 2: 2 workers and 2 gamergates, experiment 3: 2 workers and 2 gamergates), experiment was regressed out along with nUMI using a linear model during the scaling step. After normalizing and scaling, variable genes were found using the *FindVariableGenes* function in Seurat with default parameters except x.low.cutoff=0.0125, x.high.cutoff=3,y.cutoff=0.5. 1,447 variable genes were detected using this method.

Clustering and visualization of the data were accomplished using Seurat’s linear dimension reduction (PCA) followed by construction of a tSNE. Principal components selected for use in the tSNE were chosen by running a JackStraw resampling test and choosing all components until a component had a *P*-value > 0.05. Cells were clustered using a resolution of 1.

For reclustering of neurons and glia, cells were selected from full clustering by average cluster expression of neuron markers and glia markers. Reclustering was performed as above, but with a resolution of 0.3 for glia and 0.6 for neurons. Reclustering the glia resulted in one cluster with a far lower median UMI compared to other glia cells—these cells (marked as “***” in **Fig. S3a–b**) were discarded as potential low-quality cells.

Marker genes for each cluster were determined by the Seurat FindMarkers function, with the parameters only.pos=T and min.pct=0.25.

#### Drop-seq data analysis

Drop-seq data were processed as above, but with cutoffs of 200 UMIs and a resolution of 0.6 (for full clustering and glia reclustering). For the injury experiment (**Fig. 4**), Drop-seq data for aging experiment (d5, d30, d90, and d120 workers, and d120 gamergates) and injury experiment (injury and age-matched controls) were processed together, but only control/injury cells were shown in tSNE visualization and used for the subsequent analyses.

### Reanalysis of *Drosophila* single-cell data

DGE matrices for *Drosophila* midbrain^27^ and *Drosophila* brain^28^ were downloaded from *eLife* (Figure 1 – source data 1) and GSE107451, respectively. Analysis of these DGE matrices was performed as above with modifications.

For the midbrain dataset, the minimum UMI threshold was set at 800, as in Croset et al., and replicate was regressed out along with UMI during the scaling step. The resulting 10,305 cells were clustered using a resolution of 2.5. Neurons and glia were separated using neuron and glia markers, including *nSyb* and *bdl* (**Fig. S1c**). A resolution of 2.5 was also used for re-clustering the 9,916 neurons. Very few cells were in clusters marked by glia-specific genes (in total, 373 glia); as in Croset et al., we were unable to recover known glia clusters other than astrocytes upon reclustering glia.

For the full brain data set, the minimum UMI threshold was 500 UMIs, yielding 56,875 cells, similar to the 56,902 cells analyzed in Davie et al. Cells were clustered with a resolution of 2.0, and neurons and glia were separated as above (**Fig. S1c**), resulting in 46,944 neurons and 3,614 glia (comparable to the 3,600 glia reported in Davie et al.). Neurons were re-clustered with a resolution of 0.6, while glia were re-clustered at a resolution of 0.2.

### Pseudobulk expression measurements by sample and cluster

For cluster-level pseudobulk expression analyses (e.g. **Fig. 1C**), the number of UMI in for each gene from all cells in the cluster from each sample were added together and normalized by the total number of UMI detected in cells from that sample and cluster.

### Subsampling of single-cell data

To compare *Harpegnathos* and *Drosophila* on equal footing from a technology perspective, we equalized the number of cells in the datasets while matching the UMI distribution. To accomplish this, we first reduced each data set to the same number of cells. We then ordered the cells by number of UMI, and for cells at each index sampled the UMI to be equal to the same number as the cell at that index with the lowest number of UMI. We then re-normalized and scaled the data as described above.

### Comparison of single-cell to bulk sequencing

Data from bulk sequencing of *Harpegnathos* non-visual brains from 11 workers and 12 gamergates at d120^19^ were downloaded from GSE83807 and aligned to the Hsal_v8.5 reference. Differentially expressed genes were called using DESeq2^122^. For the single-cell sequencing data, DGE matrices were made using the Drop-seq tools v2.0.0, developed by the McCarroll Lab (https://github.com/broadinstitute/Drop-seq), but using a cutoff of 1 gene per cell. This resulted in DGE matrices containing all information from the single-cell data (every read mapped to the exon of a gene, including from cells not used in clustering). The fold-change between worker and gamergate of the top 100 differentially expressed genes in bulk sequencing was compared between bulk and single-cell data.

### Transcription factors

Homologs for all *Drosophila* genes in the “transcription factors” gene group (FBgg0000745) from FlyBase^123^ were considered for analysis. The pseudobulk expression from each cluster (see above) for each transcription factor detected in single-sequencing was compared between neurons and glia to determine transcription factors specific for one of the cell types. All transcription factors were used to hierarchically cluster neuron and glia clusters, but only those with a | log fold-change | > 1 were included in the heatmap generated by pheatmap (scaled by row) in **Fig. 1e**.

### Neprilysin family phylogenetic analysis

Protein sequences for all genes from seven insects (*Harpegnathos saltator*, *Camponotus floridanus*, *Apis mellifera*, *Nasonia vitripennis*, *Drosophila melanogaster*, *Aedes aegypti*, and *Tribolium castaenum*) containing the M13 neprilysin domain were downloaded and a multiple sequence alignment was performed using msa^124^. Using phangorn^125^ on a test set, the model with the highest log-likelihood was found to be the LG model, which was used to make a distance matrix between each sample. UPGMA clustering was calculated using these distances.

### GO analysis

An in-house script (available upon request) was used for GO enrichment analysis. GO terms were assigned to *Harpegnathos* genes based off of GO terms assigned to their *Drosophila* and/or human homologs. For each ontology, enrichment for a gene set was calculated using the number of genes annotated with a particular GO terms compared to genes not in the set. The universe of genes considered included only genes detected in the relevant cell subset (e.g. only neurons or only glia). Gene sets for neuron clusters or astrocyte clusters consisted of marker genes determined per cluster by Seurat (see above). GO analyses from the “biological process” category were visualized in heatmaps of –log_10_(*P*-values) for each term in each cluster using pheatmap, scaled by row, to determine if GO terms were broadly enriched or specific for certain clusters. Only terms with a Benjamini-Hochberg adjusted *P*-value of < 0.1 were visualized on heatmaps.

### Analysis of published D. quadriceps, C. planatus, and P. canadensis RNA-seq

For RNA-seq from other Hymenoptera, homologs were identified using BLAST. Protein models from NCBI annotations for *D. quadriceps* (ASM131382v1) and *P. canadensis* (ASM13138v1), as well as protein models from *C. floridanus* (Cflo_v7.5^23^; due to the lack of genome and annotation for *C. planatus* but their very close evolutionary relationship, as in ref^93^) were queried against a BLASTp database created from *Harpegnathos* protein models (Hsal_v8.5^23^). Homologs were defined as the gene with the best protein model match, with a e-value threshold of 10^-5^. RNA-seq FASTQ files were downloaded from GEO (*D. quadriceps* and *P. canadensis*, GSE59525; *C. planatus*, PRJNA472392) and were mapped against the assemblies above using STAR v2.4.1d. Gene-level counts and RPKMs were calculated using GenomicRanges.

### Bulk RNA-seq analyses in *Drosophila* brains

RNA-seq reads were mapped to the *Drosophila melanogaster* assembly BDGP6 pre-indexed with transcript models from Ensembl 87 using STAR 2.5.0b^126^ with default parameters except -- alignIntronMax set to 10,000. Aligned reads were assigned to gene models using the summarizeOverlaps function of the GenomicRanges R package. Reads per kilobase per million (RPKMs) were calculated with a slight modification, whereby only reads assigned to annotated protein-coding genes were used in the denominator, to minimize batch variability due to different amounts of contaminant ribosomal RNA. Differential expression was determined using the DESeq2 package^122^ and visualized by conventional MA plots using the log_2_(fold-change) calculated from RPKMs. Because Davie et al. identified two slightly different clusters of ensheathing glia cells^28^ (referred to as ens-A and ens-B), we only considered markers those genes expressed in both.

### Other data sets used

Bulk sequencing data from *Harpegnathos* workers and gamergates at day 120^19^ were downloaded from GSE83798. Single-cell RNA-seq from *Drosophila* brains was downloaded from supplementary material from^27^, and GSE107451^28^. Bulk RNA-seq data for *D. quadriceps* and *P. canadensis* were downloaded from GSE59525^92^. Bulk RNA-seq data for *C. planatus* were downloaded from PRJNA472392^93^.

## Data availability

RNA sequencing data generated for this study have been deposited in the NCBI GEO with accession number GSE135513. Sequencing data will remain private during peer review and will be released upon publication.

## ACKNOWLEDGMENTS

The authors thank Tim Christopher and Cristina Brady for technical support, Eduardo Torre for help setting up Drop-seq, and Julianna Bozler, Balint Kacsoh, Karl Glastad, Linyang Yu, Danny Reinberg, Shelley Berger, and Claude Desplan for comments on the manuscript. R.B. was supported in part by the Searle Scholars Program (15-SSP-102) and a NIH New Innovator Award DP2MH107055. This project was supported by a New Initiative Research Grant from the Charles E. Kaufman Foundation to R.B. and A.R. (KA2016-85223). A.R. was also supported by a NIH Center for Photogenomics grant (RM1 HG007743) and an NIH Transformative Research Award (R01GM137425).

## SUPPLEMENTAL FIGURE LEGENDS

**Figure S1.**
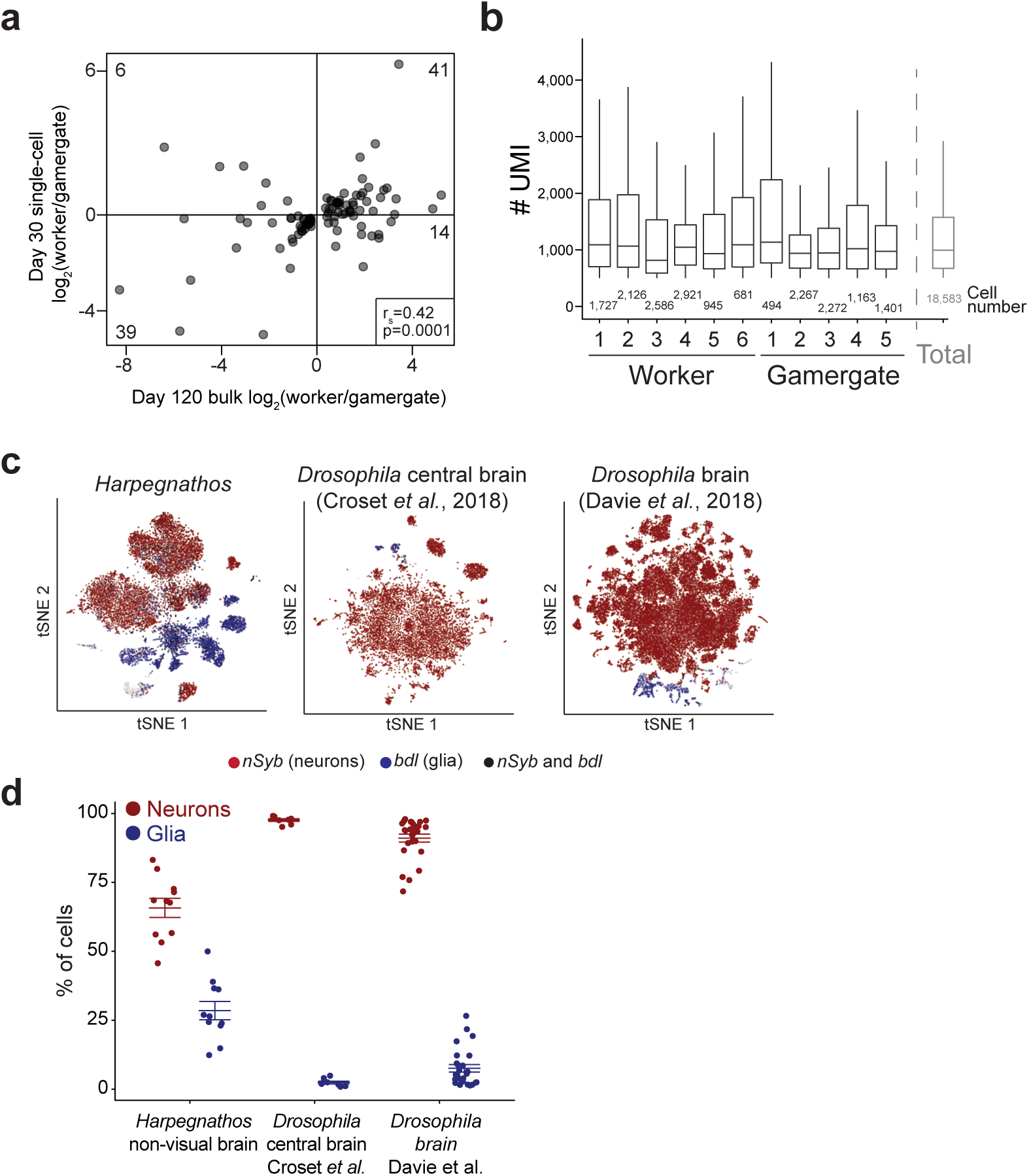
Single-cell dataset features; frequency of neurons and glia. **a**, Correlation scatter plot for previously identified caste-specific genes in *Harpegnathos* workers and gamergates comparing expression levels in bulk RNA-seq at day 120 of the transition (reanalyzed from Gospocic et al. 2017) and the 10x Genomics single-cell RNA-seq dataset obtained at day 30 for this study. Numbers of genes in each quadrant are shown. **b**, Boxplots showing the distribution of UMIs in all cells included in clustering for the separate replicates (black) and the pooled data (gray) used for clustering and tSNE visualization in Fig. 1. The number of cells per sample are indicated. **c**, Expression patterns plotted over tSNE for *nSyb* (neurons, red) and *bdl* (glia, blue) expression in cells from the current study in *Harpegnathos* (left), the *Drosophila* central brain (middle; Croset et al., 2018), and the *Drosophila* whole brain (right; Davie et al., 2018). Cells with expression of both markers are depicted in black. **d**, Relative frequency (as % of total cells) of neurons (red) or glia (blue) in *Harpegnathos*, *Drosophila* central brain (Croset et al., 2018), and *Drosophila* whole brain (Davie et. al, 2018). Horizontal bars indicate means ± SEM.

**Figure S2.**
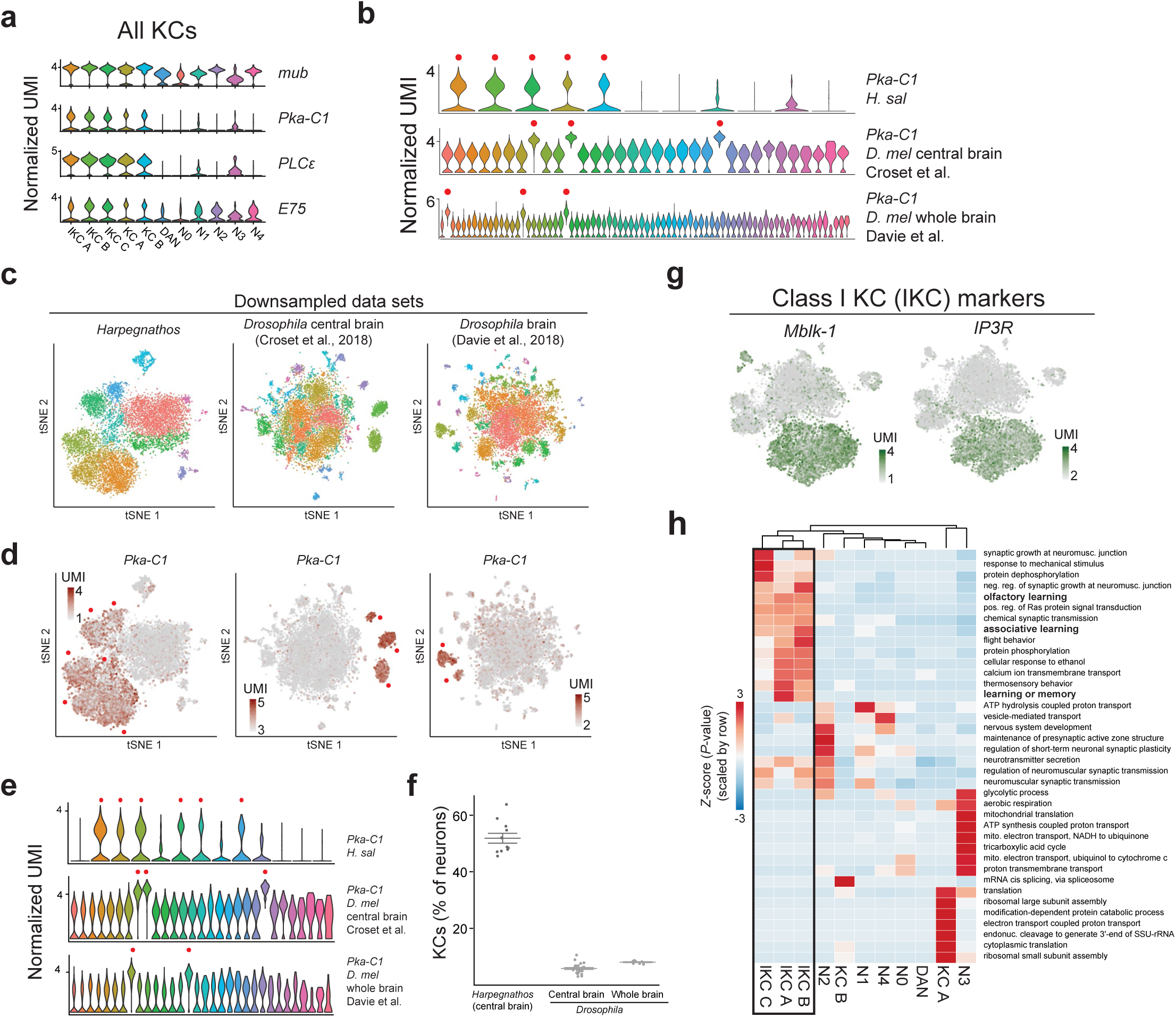
Mushroom body neurons in *Harpegnathos* and *Drosophila*. **a**, Violin plots showing expression (normalized UMIs) of mushroom body and KC markers in all neuronal clusters. **b**, Violin plots showing expression (normalized UMIs) for *Pka-C1* in *Harpegnathos* and *Drosophila* central brain and whole brain for all neuronal clusters. Red dots indicate clusters with elevated expression included in the quantifications. **c**, Visualization of the indicated datasets by tSNE after downsampling to the same number of cells and UMI/cell distribution. **d**, Heatmap plotted over downsampled tSNE showing the expression (normalized UMIs) of *Pka-C1* in *Harpegnathos* and *Drosophila* neurons. **e**, Violin plots showing expression (normalized UMIs) for *Pka-C1* in *Harpegnathos* and *Drosophila* central brain and whole brain for all neuronal clusters after downsampling. Red dots indicate clusters with elevated expression included in the quantifications. **f**, Relative frequency of KCs as determined by percentage of neurons in clusters that express *Pka-C1* in *Harpegnathos* brains and in the two *Drosophila* single-cell RNA-seq datasets after downsampling. Horizontal bars indicate means ± SEM. **g**, Heatmap plotted over neuronal tSNE for two IKC markers previously described in the honey bee. **h**, Clustered heatmap of the *P*-values for the enrichment of GO terms (rows) associated with genes specifically expressed in each cluster (columns). The heatmap contains all biological process GO terms with an adjusted *P*-value < 0.1 for at least one cluster. The color scale represents -log_10_(*P*-value) scaled by row. Terms enriched in genes expressed specifically in the lKC A, B, and C clusters are indicated with a black box. Terms mentioned in the text are bolded.

**Figure S3.**
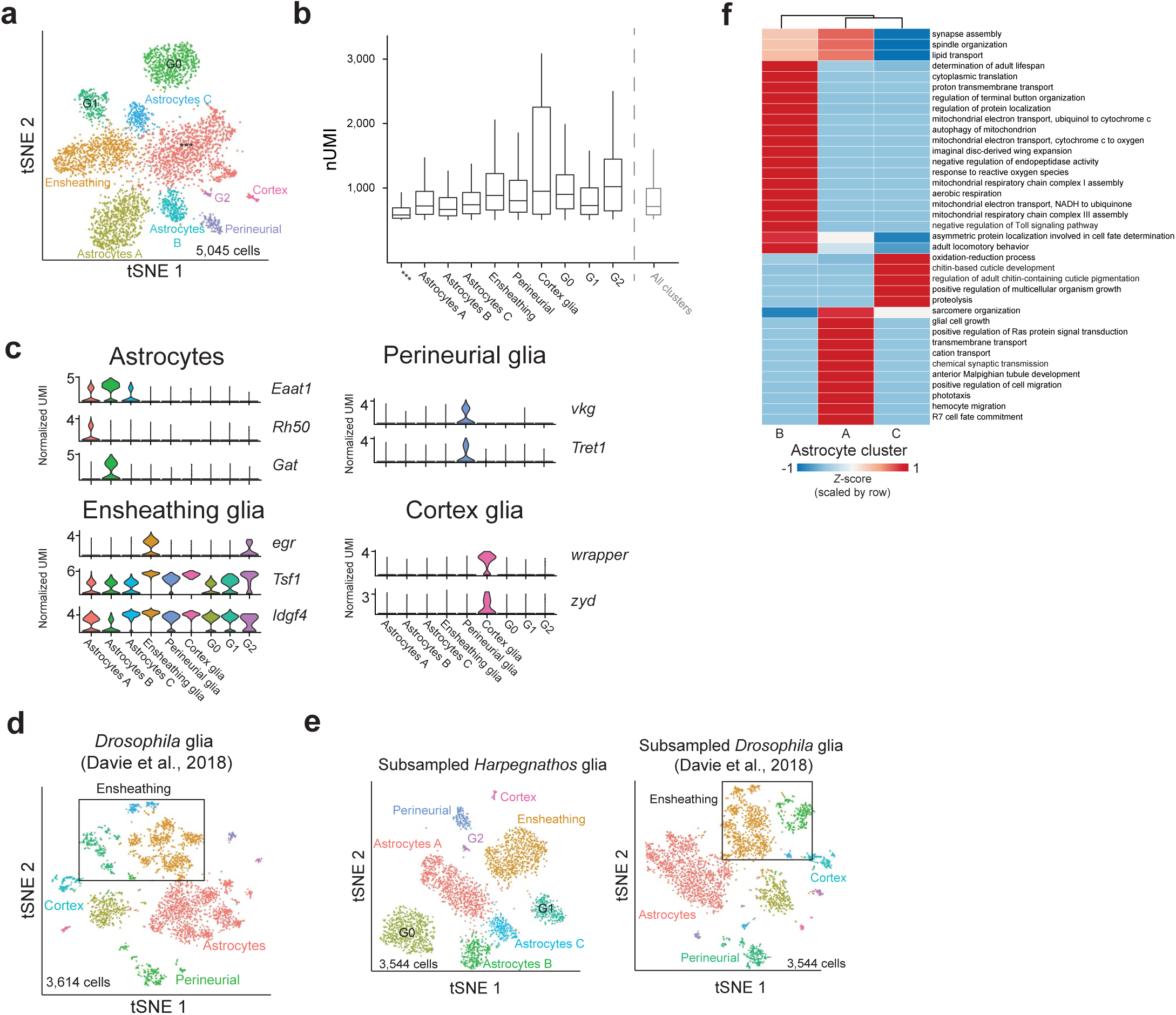
Glia in *Harpegnathos* and *Drosophila*. **a**, Annotated tSNE visualization of glia-only reclustering including the low-quality cluster, labeled “***”. **b**, Boxplot showing the distribution of UMIs in all clusters from (a) including the low quality (“***”) cluster. **c**, Violin plots showing expression (normalized UMIs) in all glia clusters for marker genes of the indicated glia subtypes. **d**, Annotated tSNE visualization of glia-only reclustering of single-cell transcriptomes from the *Drosophila* brain (Davie et al., 2018). **e**, Annotated tSNE visualization of *Harpegnathos* and *Drosophila* (Davie et al., 2018) glia after downsampling to obtain the same number of cells and UMI/cell distributions. **f**, Clustered heatmap of the *P*-values for the enrichment of GO terms (rows) associated with genes specifically expressed in each of the three *Harpegnathos* astrocyte clusters (columns). The heatmap contains all biological process GO terms with an adjusted *P*-value < 0.1 for at least one cluster. Colors represent –log_10_(*P*-value) scaled by row.

**Figure S4.**
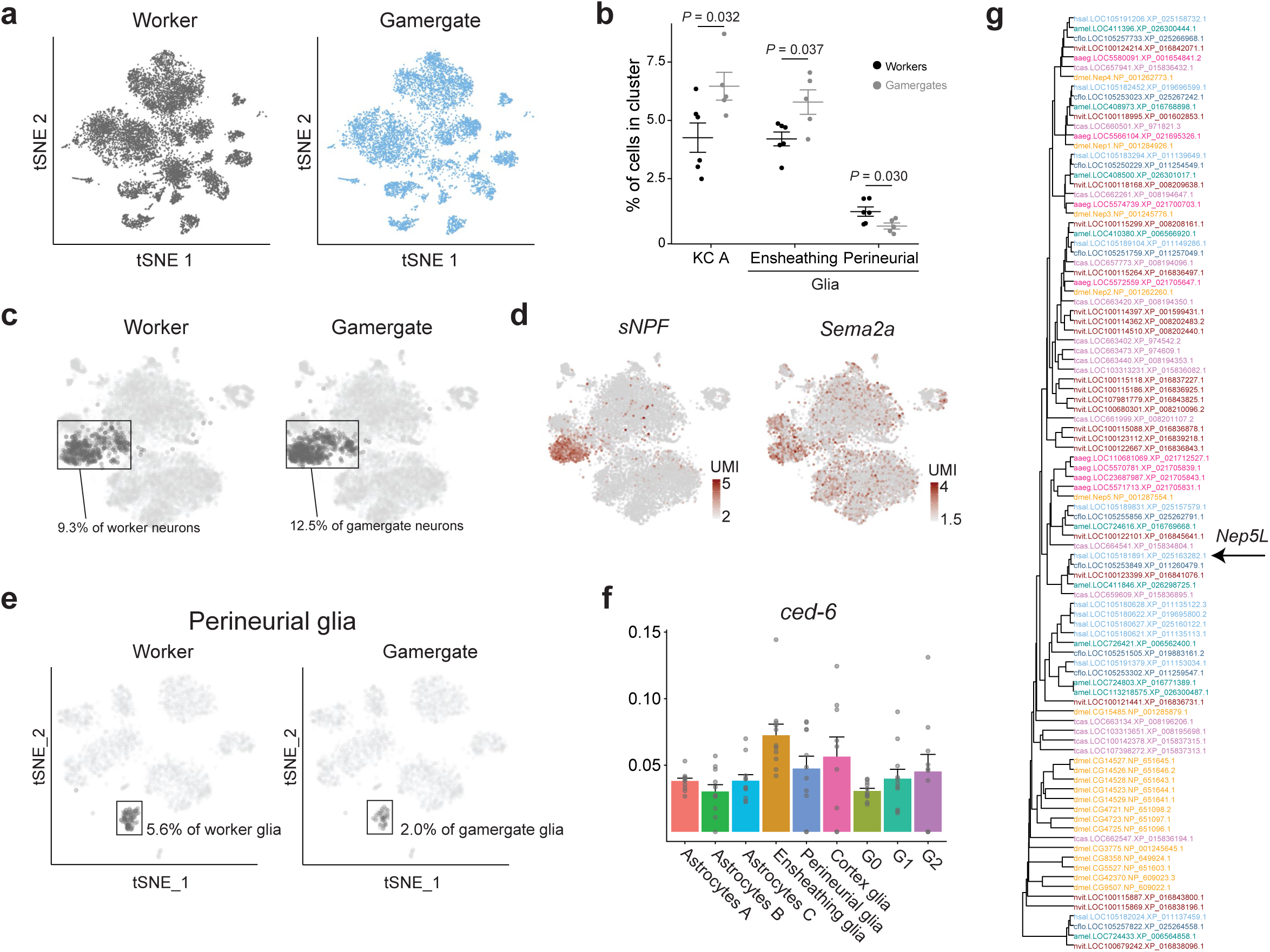
Additional analyses on caste-specific single-cell RNA-seq. **a**, Global tSNE visualization for single cells from all worker (left) and gamergate (right) samples. Worker and gamergate datasets were downsampled to include the same number of total cells for comparison. **b**, Relative frequency (as % of total cells) of cells in three caste-biased clusters. Horizontal bars indicate mean ± SEM. *P*-values are from two-tailed Student’s *t* tests. **c**, Visualization and quantification of worker (left) and gamergate (right) contributions to the KC A cluster in the reclustered tSNE for neurons only. Worker and gamergate datasets were downsampled to include the same number of total cells for comparison. **d**, Heatmaps plotted over neuronal tSNE indicating the single-cell expression levels for the indicated genes as normalized UMIs. **e**, Visualization and quantification of worker (left) and gamergate (right) contributions to the perineurial glia cluster in the reclustered tSNE for glia only. Worker and gamergate datasets were downsampled to include the same number of total cells for comparison. **f**, Expression levels (% of cluster UMIs) across glia subsets for *Ced6*. Bars indicate means + SEM. **g**, Phylogenetic tree of all genes containing an M13 neprilysin domain in 7 insects: *Harpegnathos saltator* (hsal), *Camponotus floridanus* (cflo), *Apis mellifera* (amel), *Nasonia vitripennis* (nvit), *Drosophila melanogaster* (dmel), *Aedes aegypti* (aaeg), and *Tribolium castaneum* (tcas).

**Figure S5.**
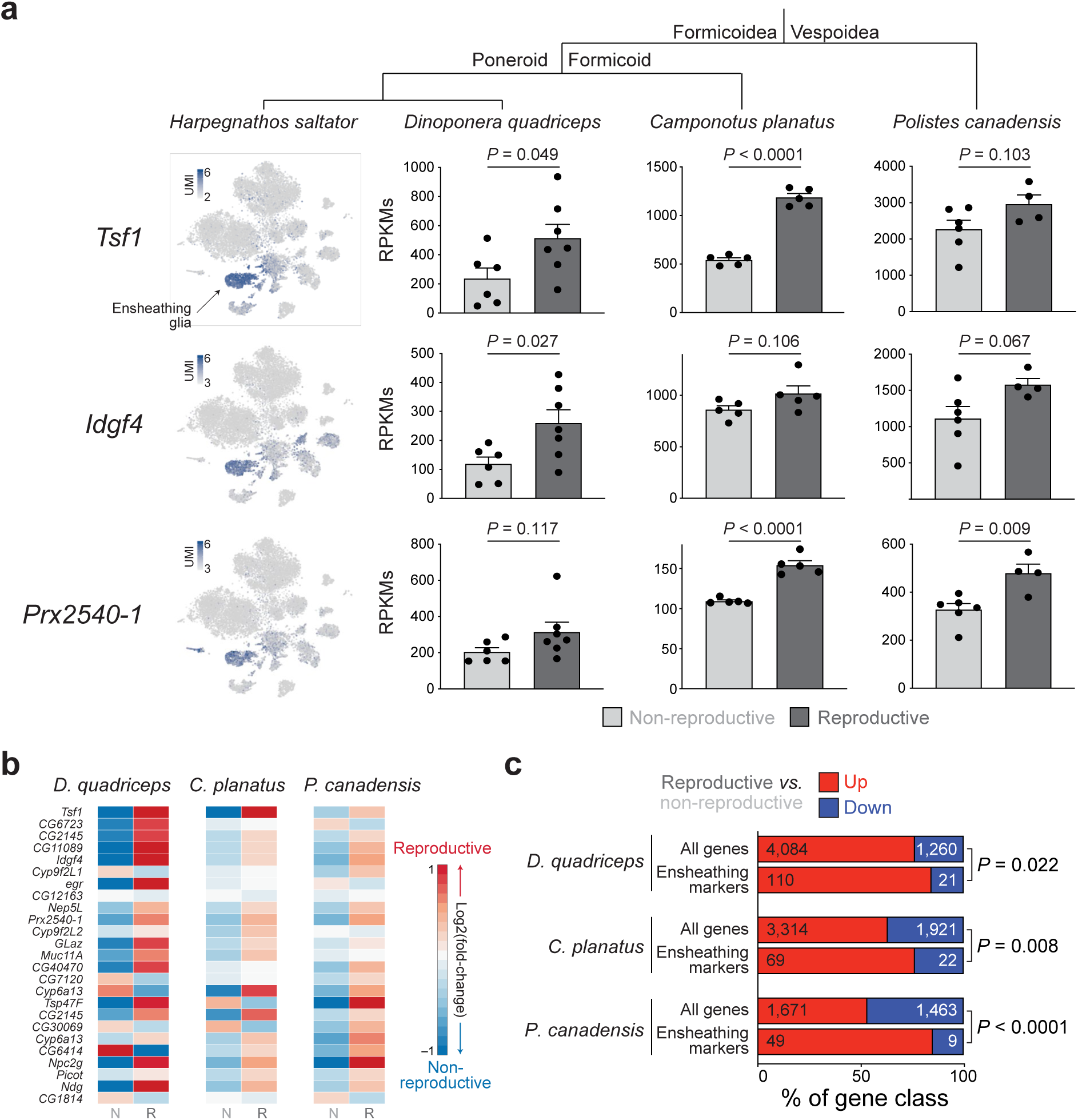
Caste-specific expression of ensheathing glia markers in other Hymenoptera. **a**, Expression patterns for three top ensheathing glia markers in the single-cell RNA-seq from *Harpegnathos* brain (left) and three published RNA-seq datasets from the brains of reproductive and non-reproductive individuals from the ponerine ant *Dinoponera quadriceps*, the ant *Camponotus planatus*, and the red paper wasp *Polistes canadensis*^92, 93^. Expression levels are plotted as reads per kilobase per million (RPKMs). Bars show means + SEM. *P*-values are from Student’s *t*-tests. **b**, Heatmap of relative expression changes in the same RNA-seq datasets utilized in (a) for the homologs of the top 25 marker genes for ensheathing glia in *Harpegnathos*. Data and is expressed as relative log_2_(fold-change) in RPKMs for reproductive (left) and non-reproductive (right) compared to the other caste. Genes with higher expression in brains of reproductive individuals are in red and genes with higher expression in brains of non-reproductive individuals are in blue. **c**, Bars show the ratio between the number of genes upregulated or downregulated (by at least 25%) in reproductive vs. non-reproductive individuals when considering all genes (gray bars) or only the homologs to the 213 genes that mark ensheathing glia in *Harpegnathos* (see Table S2). *P*-values are from Fisher’s tests.

**Figure S6.**
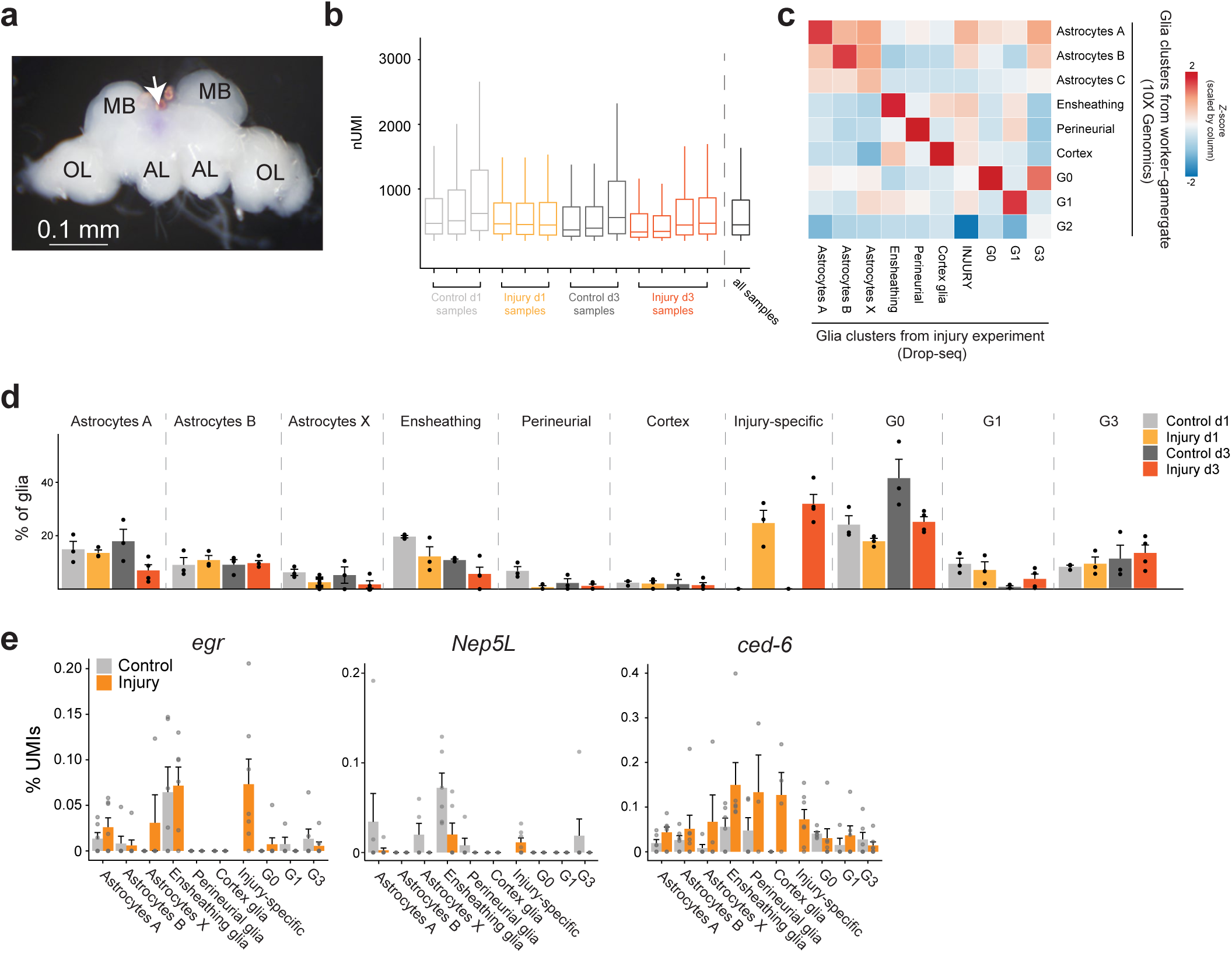
Cellular response to brain injury. **a**, Microphotograph of a dissected *Harpegnathos* brain after a stabbing injury delivered by puncturing with a needle. The site of stabbing injury was visualized with a blue dye (arrow). AL, antennal lobe; MB, mushroom body; OL, optic lobe. **b**, Boxplots showing the distribution of UMIs in all clusters for the injury Drop-seq experiment. **c**, Heatmap showing the *z*-score for the pairwise Pearson correlation between collapsed transcriptomes (pseudo-bulk analysis) of glia clusters from the day 30 worker vs. gamergates comparison (10x Genomics, Fig. 2g) and from the injury experiment (Drop-seq, Fig. 4b), considering only variable genes that were utilized to define the clusters by Seurat. **d**, Relative frequency (as % of all glia) of each cluster in control and injury day 1 and day 3. Bars represent means + SEM. **e**, Expression levels (as % of cluster UMIs) for ensheathing glia markers *egr* and *Nep5L* and the phagocytosis gene *ced-6*. Bars indicate means + SEM.

**Figure S7.**
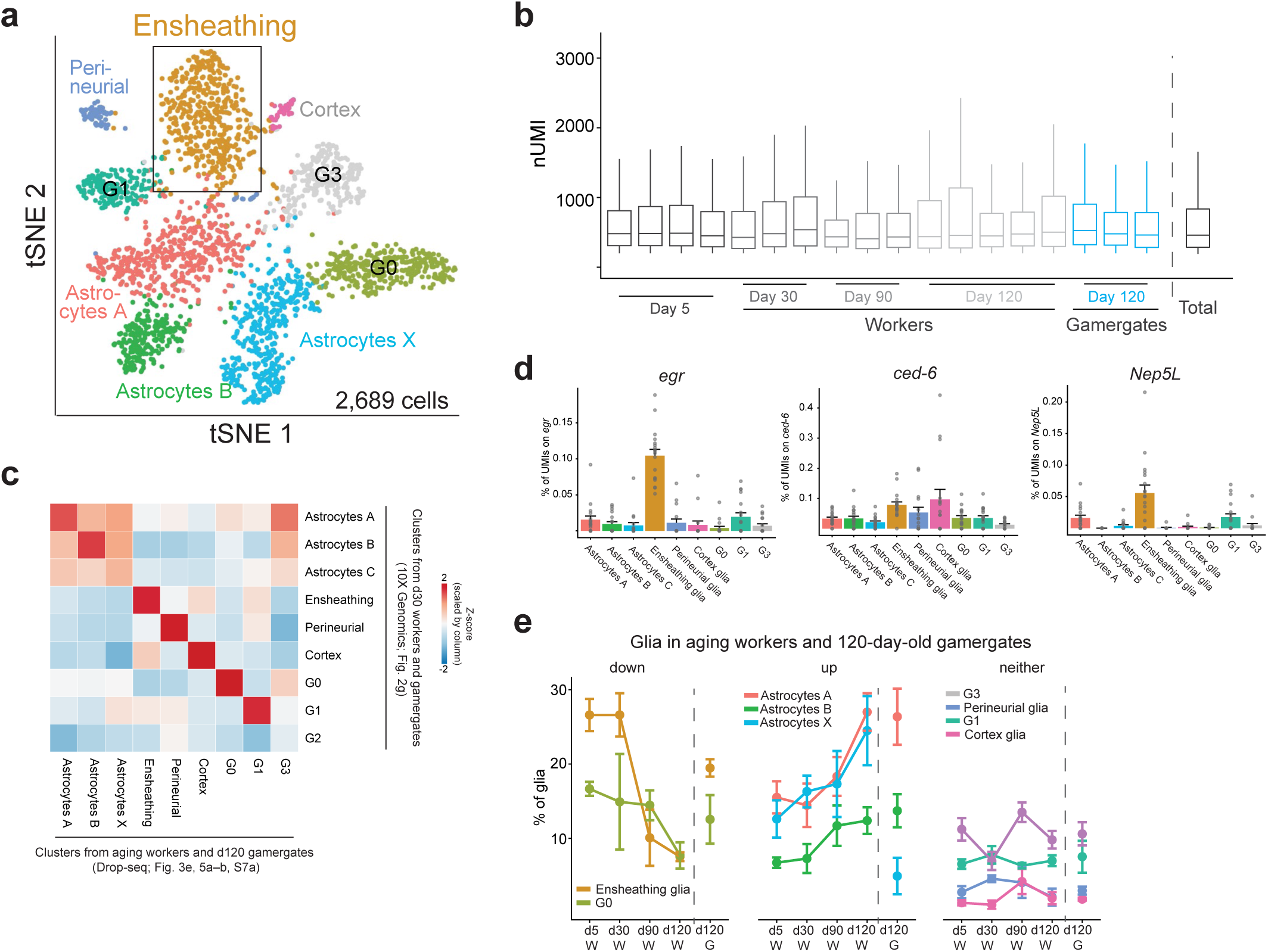
Aging-related changes in cellular composition in *Harpegnathos* brains. **a**, Annotated tSNE visualization of glia-only reclustering from Drop-seq performed on brains from workers of different ages (day 5, *n =* 4; day 30, *n =* 3; day 90, *n =* 3, day 120, *n =* 5). Drop-seq data from the 120-day-old gamergates shown in Fig. 3e were clustered with these datasets. The position of ensheathing glia in the tSNE is indicated by the box. **b**, Boxplots showing the distribution of UMIs for the brain Drop-seq datasets from aging workers (Fig. 5a–b) and day 120 gamergates (Fig. 3e). **c**, Heatmap showing the *Z*-score for the pairwise Pearson correlation between collapsed transcriptomes (pseudo-bulk analysis) of glia clusters from the day 30 worker vs. gamergates comparison (10x Genomics, Fig. 2g) and from the aging experiment (Drop-seq, Fig. S7a), considering only variable genes that were utilized to define the clusters by Seurat. **d**, Expression levels (% of cluster UMIs) across glia subsets for *egr*, *ced-6*, and *Nep5L*. Bars indicate means + SEM. **e**, Relative frequencies (as % of total glia) of all glia clusters in all samples from the aging Drop-seq experiment. Circles indicate the means, error bars represent ± SEM.

**Figure S8.**
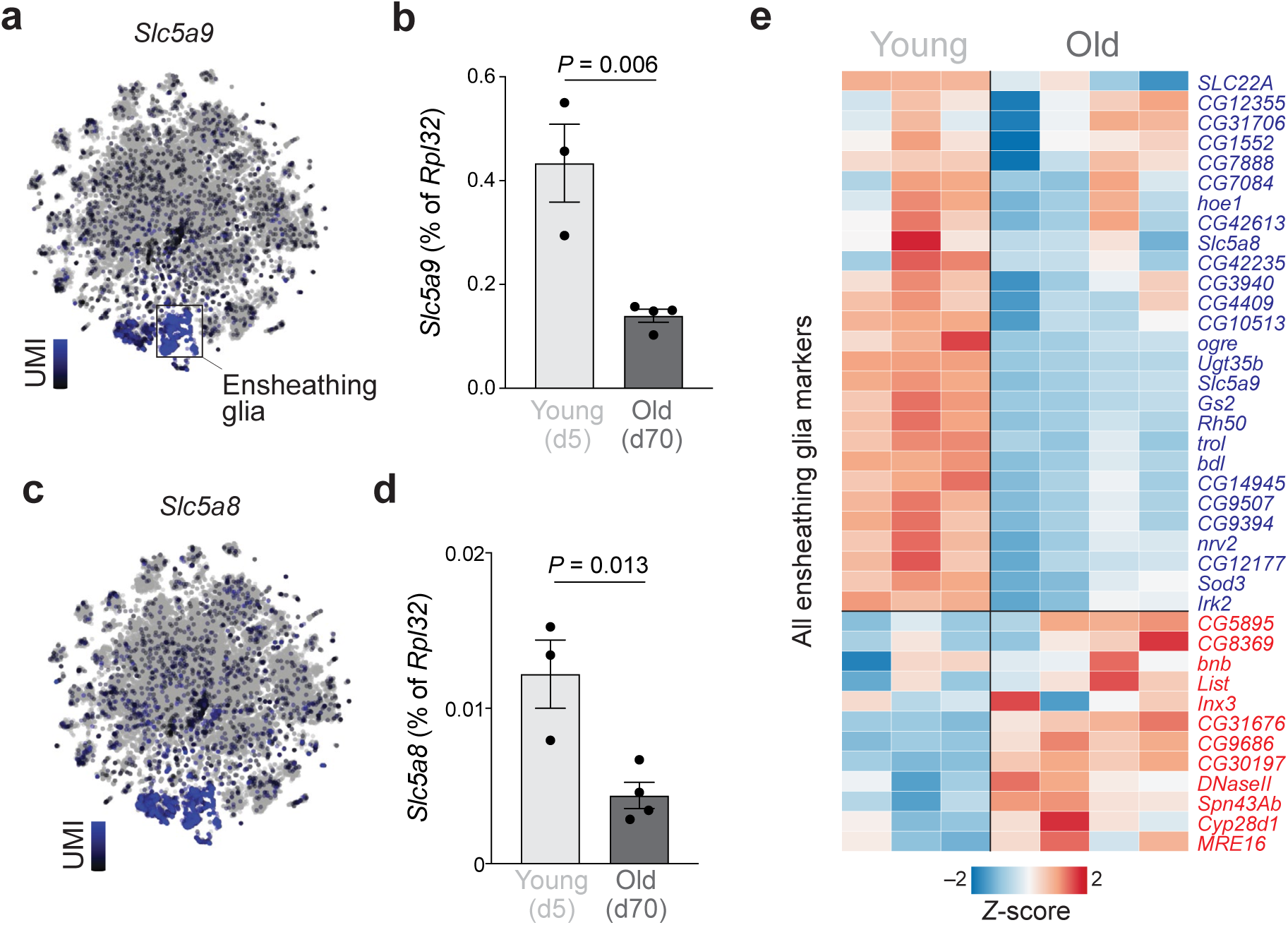
Aging-associated decrease of ensheathing glia marker expression in *Drosophila*. **a**, Heatmap of UMI levels per cell plotted over tSNE for *Scl5a9* (*CG9657*) in *Drosophila* brains. Single-cell RNA-seq data are from Davie et al.^28^. **b**, Abundance of *Slc5A9* mRNA as determined by RT-qPCR (normalized to *Rpl32*) from dissected brains of day 5 (young) and day 70 (old) *Drosophila* females. *P*-value is from a Student’s *t*-test. **c–d**, Same as in (a–b) but for *Slc5a8* (*CG6723*). **e**, Heatmap of the expression levels (RPKMs) converted to *Z*-scores for all ensheathing glia marker genes in individual RNA-seq replicates from young (day 5) and old (day 70) *Drosophila* brains.

## SUPPLEMENTAL TABLES

Table S1. Details on cluster annotation

Table S2. Marker genes for all clusters

Table S3. *Harpegnathos* gene IDs, symbols, and corresponding *Drosophila* homologs

Table S4. Marker genes for reclustered neurons

Table S5. Marker genes for reclustered glia

Table S6. Marker genes for glia clusters in the injury Drop-seq experiment

Table S7. Marker genes for glia clusters in the aging Drop-seq experiment

Table S8. Oligonucleotide sequences

